# AGORA2: Large scale reconstruction of the microbiome highlights wide-spread drug-metabolising capacities

**DOI:** 10.1101/2020.11.09.375451

**Authors:** Almut Heinken, Geeta Acharya, Dmitry A. Ravcheev, Johannes Hertel, Malgorzata Nyga, Onyedika Emmanuel Okpala, Marcus Hogan, Stefanía Magnúsdóttir, Filippo Martinelli, German Preciat, Janaka N. Edirisinghe, Christopher S. Henry, Ronan M.T. Fleming, Ines Thiele

**Affiliations:** School of Medicine, National University of Galway, Galway, Ireland; Integrated BioBank of Luxembourg, Dudelange, Luxembourg; Department of Psychiatry and Psychotherapy, University Medicine Greifswald, Greifswald, Germany; Czech University of Life Sciences Prague, Czech Republic; Center for Molecular Medicine, University Medical Center Utrecht, Utrecht, The Netherlands; Leiden Academic Centre for Drug Research, Leiden University, Leiden, The Netherlands; Computation Institute, University of Chicago, Chicago, IL, USA; Mathematics and Computer Science Division, Argonne National Laboratory, Argonne, Illinois, USA; Division of Microbiology, National University of Galway, Galway, Ireland; APC Microbiome Ireland, Cork, Ireland

## Abstract

The human microbiome influences the efficacy and safety of a wide variety of commonly prescribed drugs, yet comprehensive systems-level approaches to interrogate drug-microbiome interactions are lacking. Here, we present a computational resource of human microbial genome-scale reconstructions, deemed AGORA2, which accounts for 7,206 strains, includes microbial drug degradation and biotransformation, and was extensively curated based on comparative genomics and literature searches. AGORA2 serves as a knowledge base for the human microbiome and as a metabolic modelling resource. We demonstrate the latter by mechanistically modelling microbial drug metabolism capabilities in single strains and pairwise models. Moreover, we predict the individual-specific drug conversion potential in a cohort of 616 colorectal cancer patients and controls. This analysis reveals that some drug activation capabilities are present in only a subset of individuals, moreover, drug conversion potential correlate with clinical parameters. Thus, AGORA2 paves the way towards personalised, predictive analysis of host-drug-microbiome interactions.

## Introduction

Trillions of microbes are inhabiting our gastro-intestinal tract, with a high species and strain diversity between individuals depending on, e.g., sex, age, geographic and ethnic origin, lifestyle, and health status^1^. These microbes, collectively called microbiota, contribute essential nutrients, such as short chain fatty acid, hormones, and neurotransmitters, to human metabolism^2^. Importantly, this host-microbiota co-metabolism also extends to drug metabolism^3^. At least 15 named microbial enzymes can metabolise over 50 commonly prescribed drugs^4^ resulting in activation, inactivation, detoxification, or re-toxification depending on the drug^3^ (Figure 1). Accordingly, human gut microbes have been shown to metabolise 176 of 271 tested drugs^5^. However, the extent, to which the different species metabolise human-targeted drugs, remains largely unknown due to the lack of large-scale analysis of the distribution of drug-metabolising enzymes in microbial genomes. Consequently, it is currently not possible to estimate differences in drug response between individuals caused by different microbiota composition. Personalised therapeutic interventions that take diet, genetics, and the microbiome into account have been proposed as a promising strategy to improve treatment efficacy^6^. However, such a systems level approach requires computational, predictive modelling^6^.

**Figure 1:**
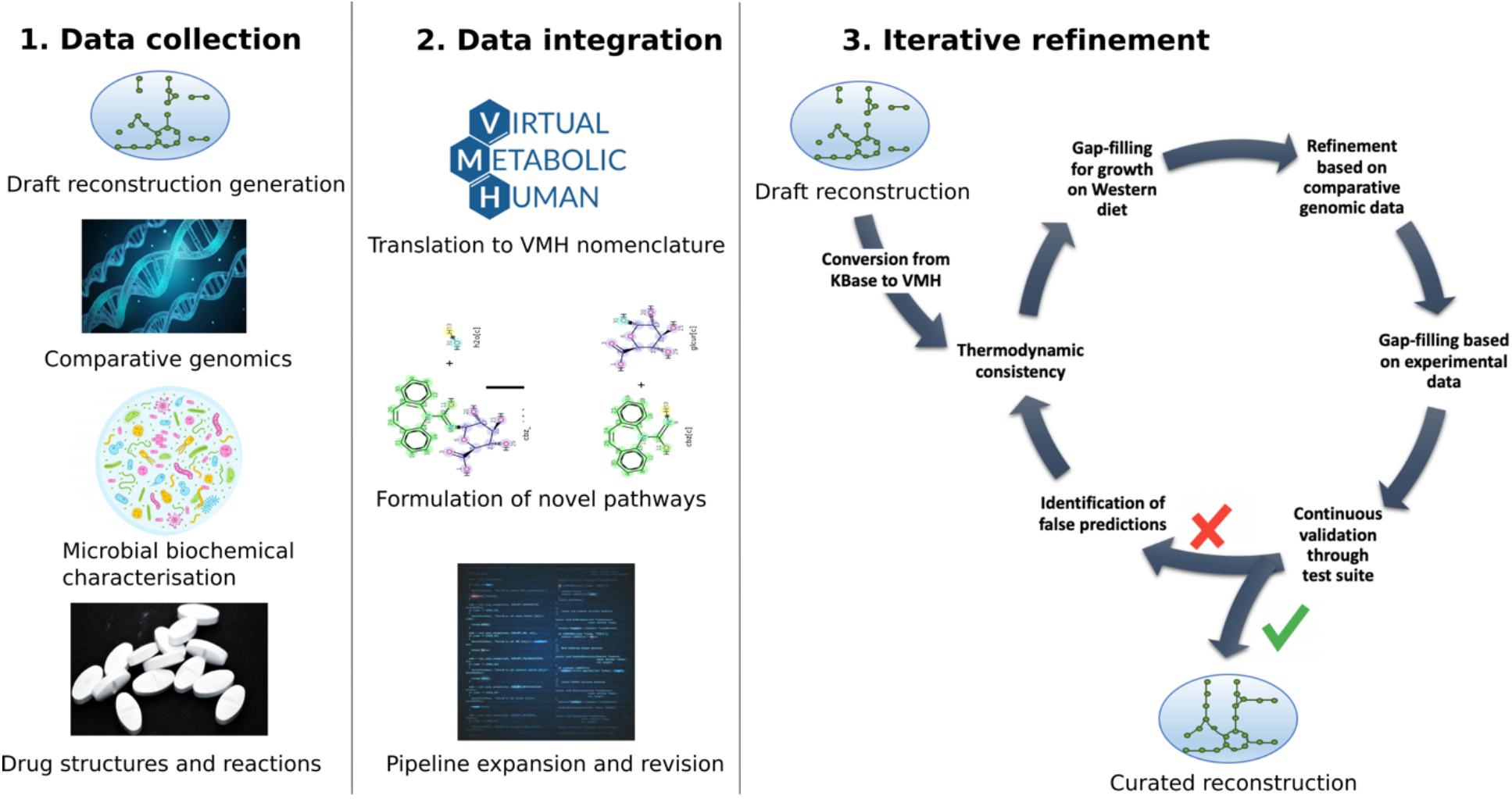
Overview of the AGORA2 pipeline. The pipeline consists of (i) collection of draft reconstructions, comparative genomic data, biochemical and physiological data, and drug structures and microbial conversion reactions, (ii) conversion of data into a MATLAB-readable format, and integration into pipeline functions, (iii) data-driven refinement of KBase draft reconstructions with continuous testing against input data and subsequent update to pipeline functions to correct for false predictions.

A mechanistic, data-driven systems approach that enables large-scale predictions of human and microbial metabolism is constraint-based reconstruction and analysis (COBRA). COBRA relies on biochemically detailed, molecule-resolved genome-scale reconstructions that are manually curated based on the available literature and represent a genomically, genetically, and biochemically structured knowledge base of the target organism^7^. These reconstructions can be converted into predictive computational models through the application of condition-specific constraints^8^, including (meta-) omics and nutritional data. Importantly, COBRA has been already successfully applied for the exploration of metabolic human-microbiome co-metabolism^9, 10^, which has been facilitated by the increasing availability of genome-scale reconstructions for human microbial species^11-13^. For instance, we have assembled a semi-automatically curated resource (AGORA1) of 773 genome-scale reconstructions of human gut microbe strains, representing 605 named species and 14 phyla^11^. To increase the microbial metabolic reconstruction coverage and capture more of the thousands of known species inhabiting humans^14^, a fast reconstruction tool, CarveMe, has been recently published^13^. Despite its many advantages, CarveMe does not account for manually refined genomic annotations and microbial drug-metabolism.

Here, we present an expansion in scope and coverage of AGORA, AGORA2, consisting of microbial reconstructions for 7,206 strains, 1,644 species, and 24 phyla. AGORA2 summarises the knowledge and experimental data obtained through extensive manual comparative genomics analyses and literature and textbook reviews. Importantly, over 5,000 AGORA2 reconstructions have been expanded by manually formulated microbial drug biotransformation and degradation reactions covering 98 drugs and 15 enzymes, which enables the prediction of drug degradation and biotransformation in a molecule-and strain-resolved manner. AGORA2 follows the quality standards developed by the systems biology research community^8, 15^ and is fully compatible with the generic^16^ and the organ-resolved, sex-specific, whole-body human metabolic reconstructions^17^. We demonstrate the use of AGORA2 for the prediction of microbial drug metabolism by single strains, pairwise combinations of microbes, and 616 personalised microbiomes. Taken together, the AGORA2 reconstructions can be used independently or together for investigating microbial metabolism and host-microbiota co-metabolism *in silico*.

## Results

### A data-driven refinement pipeline for large-scale microbial metabolic reconstructions

To build the reconstructions of the 7,206 gut microbial strains in the AGORA2 compendium, we substantially revised and expanded a previously developed data-driven reconstruction refinement workflow^11^. Overall, the reconstruction workflow consists of data collection, data integration, draft reconstruction generation and refinement, gap-filling and debugging, and iterative reconstruction curation (Figure 1). After expanding the taxonomic coverage (Figure 2a–b, Table S1, Supplemental Note 1), we generated draft reconstructions using genome sequences obtained from, e.g., the National Center for Biotechnology Information (NCBI, Table S1), and the online reconstruction tool KBase^12^. All reactions and metabolites of these draft metabolic reconstructions were translated into the Virtual Metabolic Human (VMH)^18^ name space and semi-automatically refined by including the manually collected genomic, biochemical, and phenotypic information (Figure 1). More specifically, for 5,438/7,206 (75%) genomes, we manually validated and improved the annotations of 446 gene functions across 35 metabolic subsystems using PubSEED^19^ (Table S2a-c). We performed an extensive literature search of 130 carbon sources, 30 fermentation pathways, 64 growth factors, consumption of 73 metabolites, and secretion of 51 metabolites resulting in information from 732 peer-reviewed papers and >8.000 pages of microbial reference textbooks resulting in information for 6,871/7,206 strains (95%) (Table S3a-e). For the remaining 336 strains, either no experimental data was available, or all biochemical tests reported in the literature were negative. A newly developed test suite ensured correct reconstruction structure, biochemical and thermodynamic consistency (TableS4, Supplemental Note 2). Using an unbiased quality measure, we determined the subset of flux and stoichiometrically consistent reactions^20^. The curated reconstructions had a significantly higher (p<1e-08) percentage of flux consistent reactions compared to the draft reconstructions (Figure S2) despite being larger in their metabolic content (Figure 2c). Consistently, the extensive refinement of the curated reconstruction based on genomic annotation and experimental resulted in average in the addition and removal of 489.84 (standard deviation (±): 421.10) and 111.02 (±61.96) reactions, respectively, per reconstruction (Figure S1). Note that our reconstructions represent knowledge bases, thus, if genetic or biochemical evidence exists for a gene or reaction, it will be included in the reconstruction. This approach is in contrast to other pipelines generating reconstructions containing only the flux consistent part (e.g., CarveMe ^13^, Path2Model^21^). This property allows the use of AGORA2 to rapidly identify current knowledge gaps, thereby enabling biological discovery. Moreover, we retrieved the metabolic structures for 1,838/3,533 (52%) metabolites and provide atom-atom mapping for 5,583 of the overall 7,300 (76%) enzymatic and transport reactions captured across all microbial reconstructions. Finally, biomass objective functions provided in the draft reconstructions from KBase were corrected according to gram status and reactions were placed in a periplasm compartment where appropriate (Supplemental Note 3).

**Figure 2:**
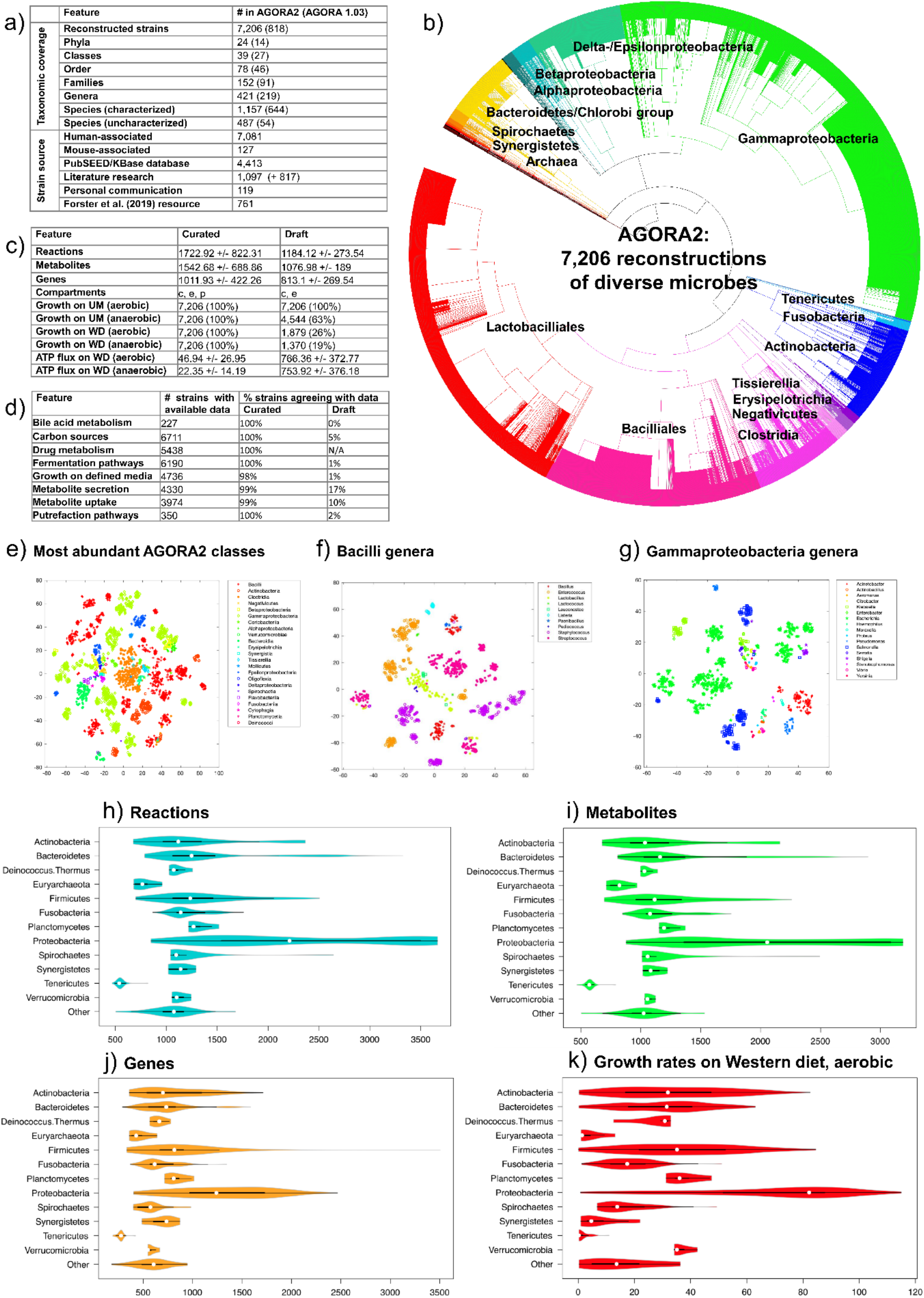
Features of AGORA2. **a)** Taxonomic coverage and sources of reconstructed strains. **b)** Taxonomic distribution of the included 7,206 strains. **c)** Features of the AGORA2 curated reconstructions and KBase draft reconstructions. c = cytosol, e = extracellular space, p= periplasm. Growth rates on Western diet (WD) and unlimited medium (UM) (Methods) are given in 1/hr. ATP production potential on WD is given in mmol/g_dry weight_/hr . **d)** Number of reconstructions with available positive findings from comparative genomics and literature, and percentage of curated and draft reconstructions agreeing with the findings for the respective organism. N/A = not applicable since pathway was absent in draft reconstructions. **e-g)** Clustering through t-distributed stochastic neighbour embedding (t-SNE)^95^ of reaction presence across all pathways per reconstruction. e) Members of the 23 largest classes by class. **f)** Members of the Bacilli class by genus. g) Members of the Gammaproteobacteria class by genus. **h-k)** Features of all AGORA2 reconstructions across phyla: **h)** Number of reactions. **i)** Number of metabolites. **j)** Number of genes, and k) Growth rate in 1/hr on aerobic WD.

The metabolic models derived from the semi-automatically curated reconstructions showed a clear improvement in predicted aerobic and anaerobic growth on unlimited medium and on a Western diet and in their agreement with experimental data over models derived from the draft reconstructions (Figure 2c, d; Supplemental Note 2). This increased predictive capability was expected as the reconstructions were curated against this data during the refinement step. Overall, AGORA2 reflects the diversity of captured strains as they clustered by class according to their reaction coverage (Figures 2e, S3a, Supplemental Note 4). Several genera in the Bacilli and Gammaproteobacteria classes formed multiple subgroups illustrating important metabolic differences between them (Figures 2f–g, S3b-c, Supplemental Note 4). Cross-phylum metabolic differences also translated to differences in predicted growth rates in Western diet (Figure 2 h–k) and in their potential to consume and secrete metabolites (Figure S4a-b). Taken together, the AGORA2 reconstructions capture current genomic and biochemical knowledge of the reconstructed microbes and can be converted into condition-specific metabolic models.

### Microbial drug metabolism guided by literature and refined genome annotations

Microbes can directly or indirectly influence drug activity and toxicity through degradation (e.g., hydrolysis) and biotransformation (e.g., reduction)^3, 4^ (Figure 3a). To account for this microbial capability, we performed an extensive, manual comparative genomic analysis for 25 drug genes, encoding for 15 enzymes shown to directly or indirectly affect drug metabolism (Table S5), their subcellular locations, and 12 genes encoding for drug-transporter (Table S2d). All 5,438 analysed strains carried at least one drug-metabolising enzyme (Figure 3a, Table S2c). As these enzymes are also involved in central metabolism, e.g., nucleoside metabolism, this high coverage was expected. We then carried out a thorough literature and database review of metabolite structures, formulas, and charges for 98 frequently prescribed drugs belonging to 10 drug groups and 32 subgroups (Figure 3b). We formulated 1,440 reactions containing 363 metabolites and added, in average, 254 reactions and 110 metabolites to the reconstructions depending on the genomic evidence (Table S6a-b). We validated, with an accuracy of 0.78 (Fisher’s exact test: p=1.50e18), the drug-metabolising predictions against independent published experimental data for 238 drug-microbe pairs (Table S7, Figure S5-S6). The 19 false positive predictions may indicate non-functional genes or regulatory mechanisms, whereas the 34 false negative predictions could be due to incompleteness of genomes or non-orthologous displacement in complete genomes. Taken together, this large-scale genome-annotation effort revealed a wide range of phyla capable of drug metabolism.

**Figure 3:**
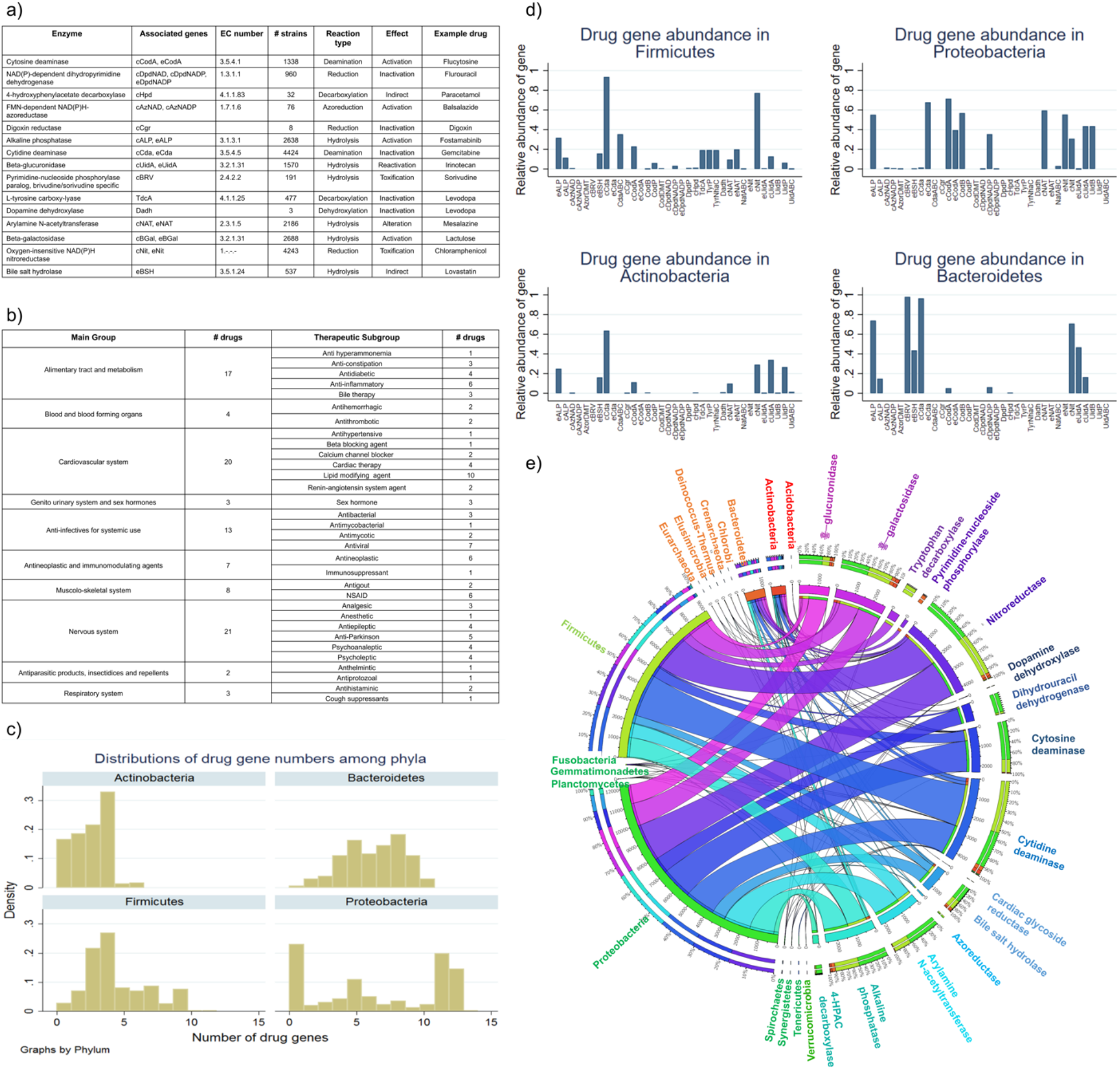
Overview of reconstructed drugs and annotated drug enzymes present in AGORA2. **a)** Description of the 15 enzymes directly or indirectly metabolising drugs that were annotated in this study. The IDs for associated genes are as given in the PubSEED subsystem and in Table S2c. ‘c’ indicates genes for cytosolic enzymes and ‘e’ indicates genes for extracellular enzymes. **b)** Description of the 98 drugs, for which microbial metabolism was reconstructed. **c)** Fraction of strains carrying each gene encoding drug enzymes or transport proteins in the four main phyla in the human microbiome. For the list of abbreviations, see Figure 3a and Table S2b. **d)** Distribution of the number of drug genes per strain for the four main phyla. **e)** Distribution of the number of strains carrying each drug enzyme over the 14 analysed phyla.

### Taxonomic distribution of drug-metabolising capabilities

We analysed the taxonomic distribution of the annotated drug and transport genes (Figure 3c–e, Table S2c). At least one strain in each of the 14 analysed phyla encoded for genes involved in drug metabolism (Figure 3e). The most widespread drug-metabolising enzymes were the central metabolic enzymes, cytidine deaminase and nitroreductase, which were found in 12 and 13 phyla, respectively (Figure S7a-b). Another central metabolic enzyme, the pyrimidine-nucleoside phosphorylase, was also widely distributed, but the monophyletic branch specific for the metabolism of brivudine and sorivudine^22^ was only found in the Bacteroidetes phylum (Figures 3d–e, S7c). Many drugs are detoxified by the liver through the addition of glucuronic acid^3^. The microbial β-glucuronidase removes glucuronic acid through hydrolysis thereby reverting the drug to its active form. This enzyme was in >99% of analysed *Escherichia coli* strains and was also widely distributed across Bacteroidetes and Firmicutes strains (Figures 3d–e, S7d), consistent with previous analyses^23^. Interestingly, *E. coli* was the species most enriched in drug metabolism with >99% of all analysed strains carrying seven to ten drug enzymes (Table S2c). The cardiac glycoside reductase and dopamine dehydroxylase could only be found in *Eggerthella lenta*, in agreement with previous reports^24, 25^. Taken together, drug-metabolising enzymes, and transporters, are widely distributed but important phyla-specific and strain specific differences exist.

### Drugs can serve as carbon, energy, and nitrogen sources and influence microbe-microbe interactions

Next, we investigated *in silico* the theoretical benefit of metabolising drugs for each microbe. Therefore, for all 5,378 reconstructions expanded with the drug reactions and for one example drug per enzyme (15 in total), we computed, using flux balance analysis^26^, the yields for ATP, carbon dioxide, pyruvate, and ammonia from 1 mmol drug/g_dry weight_/hr (Figure 4a). Of the 5378 strains, 3,828 could use carbon-containing drugs as a source of energy (ATP), carbon dioxide, and/or pyruvate (Figure 4a, Table S8). Additionally, 1,619 and 2,319 strains could use gemcitabine or 5-fluorocytosine, respectively, as a nitrogen source through deamination, and 672 strains could use taurine cleaved from taurocholate as a nitrogen source (Figure 4a, Table S6). These results suggest that the presence of drugs could alter the nutrient environment for microbes, which may have implications on interspecies growth and interactions^27^. To test this hypothesis, we simulated co-growth of 19,900 microbe-microbe pairs on three different diet compositions (Western diet^11^: alone, plus glucuronidated irinotecan (SN38G), and plus free glucuronic acid, (Table S9)). Mutualistic interactions increased from 22% to 33% when the β-glucuronidase was only in one strain and the other had only the glucuronic acid pathway (Figure 4b). In contrast, when both microbes had β-glucuronidase and the glucuronic acid degradation pathway, SN38G supplementation resulted in decrease in mutualism from 15% to 10% (Figure 4b). Taken together, the human-targeted drugs can serve *in silico* as nutrients to the microbes and may thus alter interspecies interactions.

**Figure 4:**
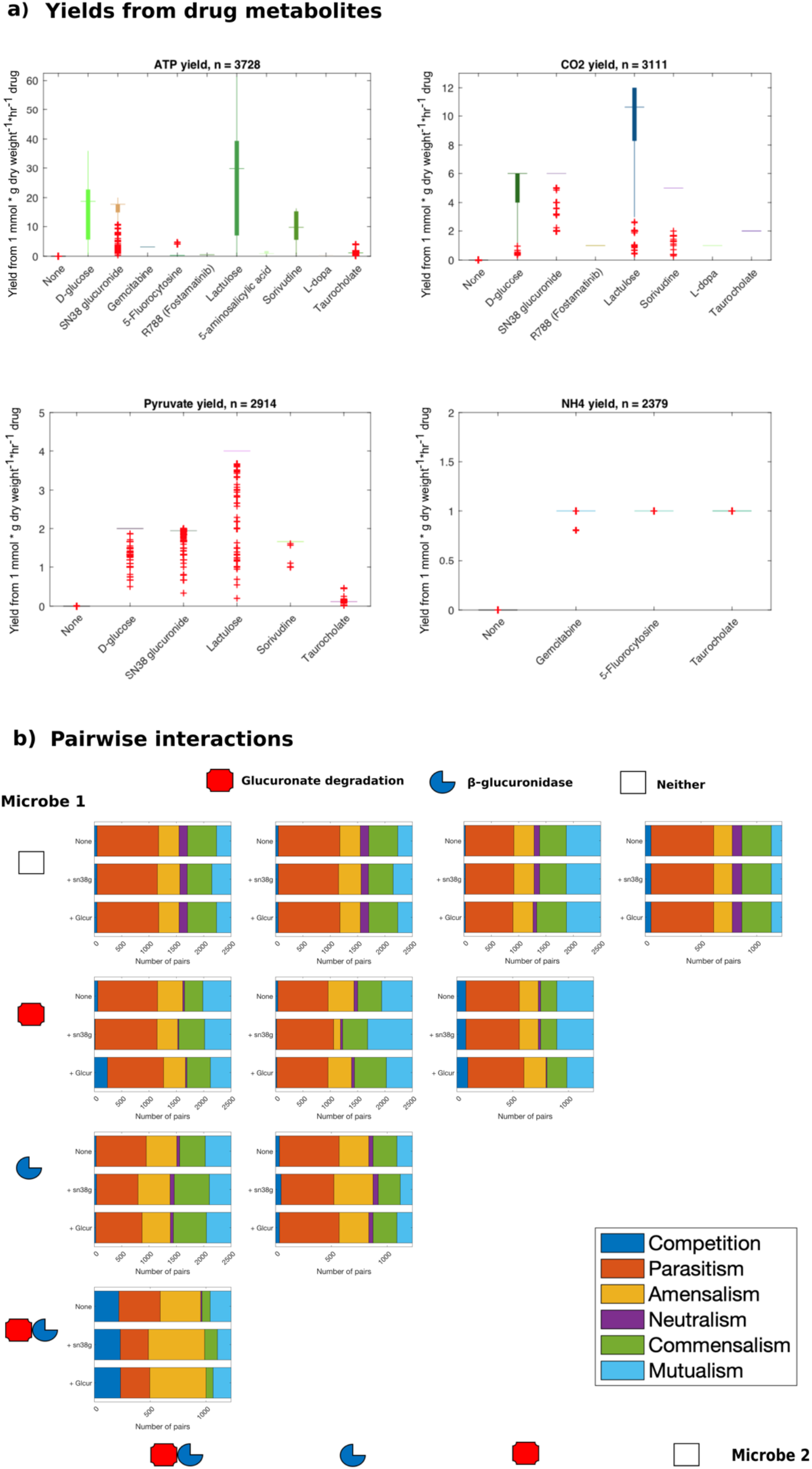
Drugs serve as nutrients for human microbes and influence microbe-microbe interactions. a) Yields from 1 mmol/g_dry weight_/hr of drugs that can serve as sources for ATP, CO_2_, pyruvate, and NH4 production. Shown are all microbes that could use at least one drug to produce the respective source. Glucose and no compound added are shown as controls. One example drug per enzyme was tested. b) Pairwise interactions between 200 microbes corresponding to 19,900 microbe-microbe pairs on an anaerobic Western diet with no additional compounds (none), SN38G added (+sn38g), and free glucuronic acid added (+glcur). Shown are percentages of the six possible interactions grouped by the characteristics of the two microbes in each pair regarding the presence of glucuronic acid degradation pathway, ß-glucuronidase, or neither.

### Community modelling reveals rare and common drug-metabolising capacities among colorectal cancer cases and healthy controls

We then addressed the important question on how the drug-metabolising capacities may differ between individuals due to different microbiota composition using a comprehensive, large-scale metagenomic data set from a Japanese cohort of 365 colorectal cancer (CRC) patients and 251 healthy controls^28^. A total of 97% of the named species could be mapped onto the AGORA2 (compared to 72% for AGORA1). For each individual, we integrated all microbial models having a non-zero abundance in the sample into one personalised microbiome model. We then computed, using flux balance analysis^26^, each individual microbiomes’ drug-metabolising potential (Figures 5, S8, Table S10). All drugs but digoxin and balsalazide could be qualitatively metabolised *in silico* by at least 95% of the microbiomes (Figure 5a, Table S10a-b), but the microbiomes’ quantitative drug-metabolising potentials varied (Figure S8). Digoxin could be metabolised by only 53% of the microbiomes presented the capacity to metabolise digoxin (Figure 5a), being strictly dependent on the presence of *Eggerthella lenta* (Figure S9). Balsalazide could be metabolised by 42% of the investigated microbiomes (Figure 5a) with a tendency of enrichment in CRC cases (Figure 5b) (odds ratio (OR)=1.39, 95%-CI=(0.99;1.96), p=0.056). Accordingly, the azoreductase was only found in 78 of 5,438 (1.4%) analysed strains (Figure 3a, Table S2c), of which only seven species were present in this cohort (Figure 6a, Supplemental Note 5, Table S11). The conversion of the prodrug 5-fluorocytosine into the active drug 5-fluorouracil and subsequent detoxification^4, 29^ was limited by the presence of certain species (Figure 6b, Supplemental Note 5, Table S11). Equally, the conversion of Parkinson’s Disease drug levodopa into m-tyramine^25^, which limits the levodopa bioavailability, was dependent on the presence of *E. lenta* in a microbiome (Figure 6c, Supplemental Note 5, Table S11). We found that diet altered only the predicted microbial community drug-metabolising potentials for 4-hydroxyphenylacetate and soriduvine highlighting putative diet-drug-metabolome interactions (Figure S10, Table S12). These examples demonstrate the added value of simulating enzymatic functions in their metabolic context rather than merely counting gene functions captured in a given microbiome sample.

**Figure 5:**
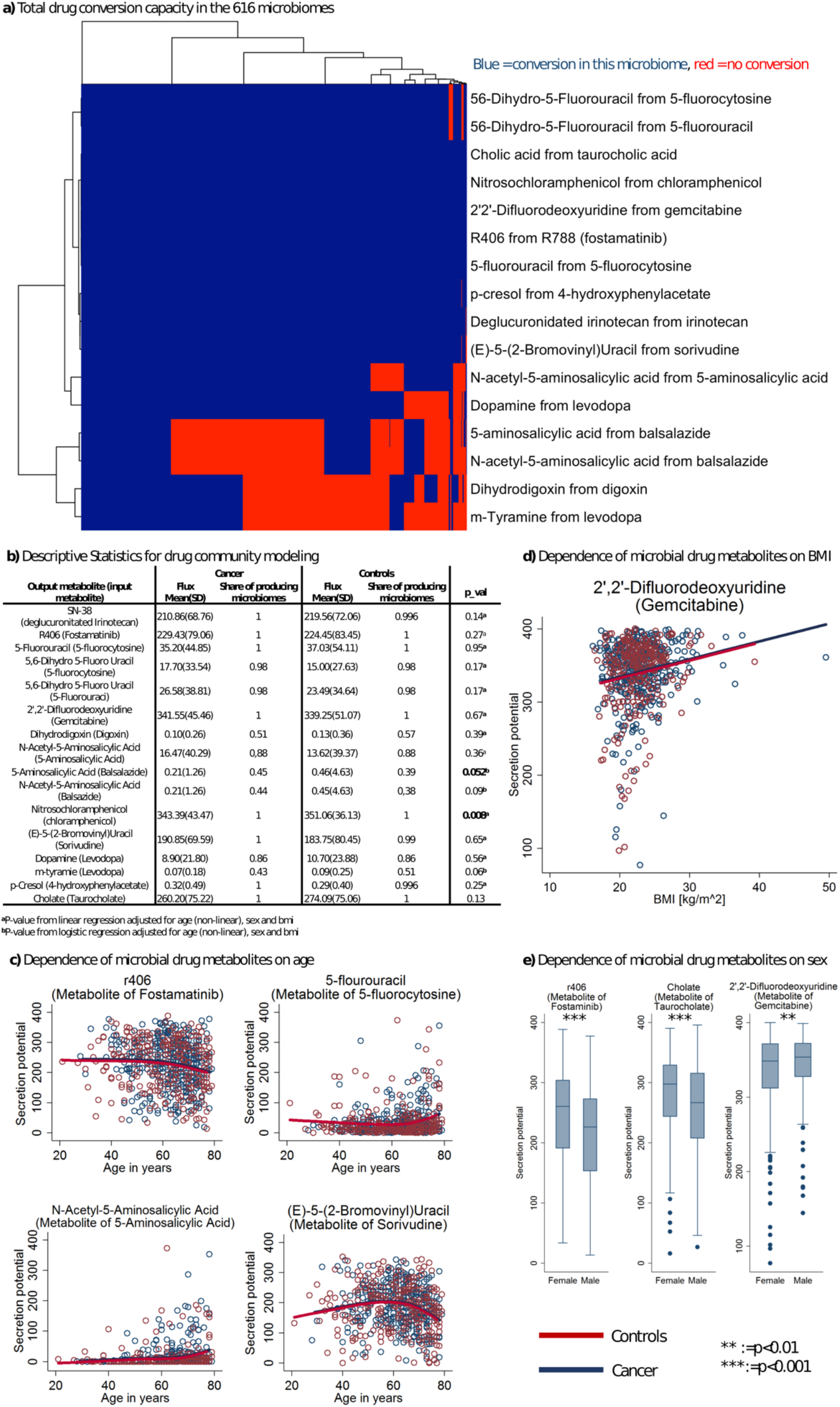
The drug conversion potential of 616 microbiomes in a Japanese cohort of 365 CRC patients and 251 controls correlates with BMI, age, sex, and CRC stage. a) Total qualitative drug conversion potential in the 616 microbiomes Blue=microbiome can convert the drug, red=no conversion. Only 53% (95%-CI=(49%;0.57%) and 42% (95%-CI=(38%;46%) of the microbiomes could metabolise digoxin and balsalazide, respectively. b) Overview on descriptive statistics for the modelled drug metabolites. c) Scatter plots (red: controls; blue cancer) of various drug metabolites in dependence on age with non-linear regression lines for cases and controls. Regression lines were estimated with restricted cubic splines. All regression models had p<0.0001 (FDR<0.05) and regression coefficients were virtually the same for cases and controls. d) Scatter plot (red: controls; blue cancer) of 2’,2’-difluorodeoxyuridine (microbial metabolite of 5-fluorocytosine) in dependence of BMI with linear regression lines for cases and controls. The slope of BMI was significant (b=2.11, 95%-CI=(1.12;3.09), p=2.88e-05, FDR<0.05) adjusted for sex and age (restricted cubic splines), but no significant differences could be found between CRC cases and controls (p=0.71). e) Box plots of 2’,2’-difluorodeoxyuridine (metabolite of gemcitabine), cholate (metabolite of taurocholate r406 (metabolite of fostamatinib) on sex. P-values were derived from linear regressions adjusted for age (restricted cubic splines). All three effects were significant after correction for multiple testing (fostamatinib: b=-31.3, 95%-CI =(−43.70;−19.05), p=7.58e-07, FDR<0.05; gemcitabine: b=12.89, 95%-CI=(4.85;20.92); cholate: b=−25.81, 95%-CI:(−37.45;−14.18), p=1.55e-05, FDR<0.05). f) Predicted share from logistic regressions of microbiomes able to produce 5-aminosalicylic acid in dependence of age (restricted cubic splines) and case-control status. Effect of age was significant corrected for multiple testing FDR<0.05.

**Figure 6:**
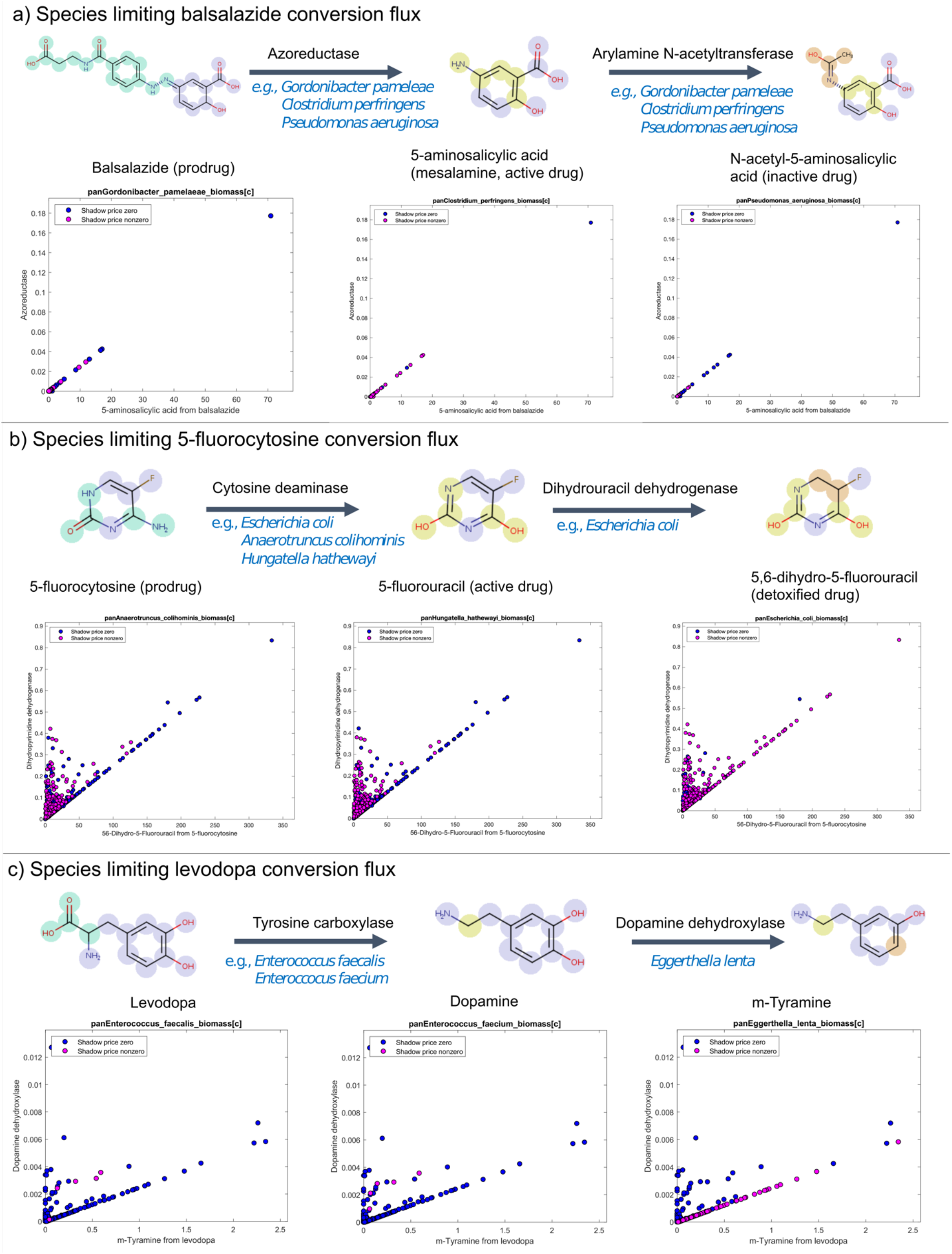
Bottlenecks limiting drug-metabolising capacity in 616 microbiomes. Non-zero shadow prices indicate that increasing the abundance of this species would increase the secretion flux of the end product of the shown enzymatic reaction in this microbiome. A shadow price of zero shows that increasing the abundance of the species would not affect secretion of the end product. a) Pathway of balsalazide azo-reduction to 5-aminosalicylic acid and subsequent acetylation to N-aetyl-5-aminosalicylic acid. b) Pathway of 5-fluorocytosine deamination to 5-fluorouracil and subsequent reduction to 5,6-dihydro-5-fluorouracil. c) Pathway of levodopa decarboxylation to dopamine and dopamine dehydroxylation to m-tyramine. In each panel, the x axis shows net secretion flux of the drug metabolite per microbiome in mmol/g_dry weight_/day and the y axis shows the relative reaction abundance per microbiome.

### Drug-metabolising capacities of microbiomes are associated with age, sex, BMI, and CRC stages

Next, we investigated the statistical association pattern of age, sex, and body mass index (BMI) to the drug-metabolising capacities of the microbiome. Predicted secretion potentials of drug metabolites were clearly associated with age (Figure 5c), although the effect sizes were small to medium. For example, the conversion of sorivudine into a toxic byproduct showed a reverse U-shaped association to age with old and young individuals having lower production potentials than middle aged persons (Figure 5c, R^2^ = 0.06, p=7.67e-08). Interestingly, women had clearly higher fostamatinib and taurocholate metabolising capability, while slightly, but significantly, lower conversion potential of the chemotherapy drug gemcitabine (Figure 5e). However, the latter effect was not significant when adjusted for BMI indicating that the lower potential in women was putatively related to the sex-differences in BMI. Indeed, conversion of gemcitabine was positively associated with BMI measures (Figure 5c), indicating that BMI acts statistically as a mediator variable.

Lastly, we investigated whether drug-metabolising capacities were associated with the CRC stage. Interestingly, conversion potential of the antibiotic chloramphenicol was clearly reduced in late stage CRC (p=0.003), which would result in increased toxicity (Figure 3a). For the other drugs, including the cancer drugs, no clear differences in drug-metabolising capacities could be observed, despite the reported enrichment in 29 species in CRC metagenomes^30^. Nonetheless, individual differences, regardless of disease status, due to distinct microbiota composition existed (Figure S8). In conclusion, AGORA2 in conjunction with metagenomic data and clinical parameters enables the investigation into the physiological and pathophysiological traits associated with drug-metabolising capacities.

## Discussion

Here, we introduced AGORA2, a resource of 7,206 genome-scale reconstructions for human-associated microbes with unprecedented coverage, scope, and curation effort. AGORA2 is freely available to the scientific community both as a knowledge base and a metabolic modelling tool^18^. AGORA2 accurately captures biochemical and physiological traits of the target organisms and includes manually refined, strain-resolved drug-metabolising capabilities.

Computational modelling of microbial consortia is increasingly recognised as a complementary method to *in vitro* and *in vivo* experiments and has the potential to generate efficiently experimentally testable hypotheses^10, 31^. Our knowledge about gut microbes remains limited and thus, any *in silico* reconstruction will be inherently incomplete and require regular updates^32^. Consequently, AGORA2 currently does not yet capture the breath of microbial drug and secondary metabolism, e.g., for plant polyphenols^33^. However, this information may be added once a strain-and enzyme-resolved understanding is obtained, as it has been done for bile acid metabolism in AGORA1^34^. When modelling microbial consortia, it is important that any reconstruction bias (e.g., set of pathways included) is consistent as it is the case for AGORA2. Moreover, as AGORA2 uses the same metabolite and reaction nomenclature^18^ as the human metabolic reconstruction^16^ and the whole-body metabolic reconstructions^17^, the microbiome-level reconstructions (and models) can be used to predict host-microbiome co-metabolism^35^, up to their potential contribution to human organ-level metabolism^17^. Available software tools^36, 37^ allow for the contextualisation of microbial and human metabolic models with omics data, thereby allowing for condition-specific and personalised modelling^17^ and for modelling of the gut (e.g.,^38^), lung^39^, tumor^40^, wound^41^, and bioremidiation^42^.

The taxonomic extension of AGORA2 (Figure 2a, b) covers all 83 named microbes in the Broad Institute-OpenBiome Microbiome Library^43^, all strains in the human gastrointestinal bacteria culture collection^44^, 99 of 100 named species in 92,143 metagenome-assembled genomes from 11,850 human gut microbiomes^45^ as well as 477 of the 573 named species (83%) in a resource of over 150,000 microbial genomes^46^. Furthermore, AGORA2 accounts also for species pre-dominantly found in non-Western populations and in disease states, which we demonstrated by investigating metagenomic samples of Japanese CRC patients and controls (Figure 5–6, Figure S8-10). AGORA2 also captures almost 500 currently uncultured and/or uncharacterised strains, and 127 mouse-associated strains (Figure 2a, Table S1). Together, this extension increases the prediction fidelity of microbiome-level models and will further broaden its application areas.

Using AGORA2, we predict that gut microbes could use a broad range of frequently prescribed drugs (Figure 3b) as energy, carbon, and nitrogen sources (Figure 4a). In fact, depending on drug and microbes, the microbe-microbe interactions were altered, which may introduce changes in the microbial ecology (which cannot be readily predicted using the COBRA approach). The predicted changes in interspecies interactions through liberation of sugars from drugs is similar to cross-feeding networks in the gut mediated by polysaccharides^47^. Competitive and mutualistic interactions, which were influenced by the presence of glucuronic acid and SN38G (Figure 4b), have been shown to destabilise microbial communities^48^. These predictions underpin increasing reports of the extent, to which gut microbes may alter the bioavailability and toxicity of human-targeted drugs^49^. The microbial communities could only be modelled on the species level, in the absence of strain-level abundances^28^. However, the well-described strain-level differences for, e.g., *E. lenta*^24^, were captured by AGORA2 (Tables S2c, S11). To harness the full potential of AGORA2 as a predictive tool, metagenomic sequencing on the strain-level would be valuable.

We reported associations between CRC patient-specific microbial drug conversion capabilities and clinical parameters, such as age and BMI (Figure 5). The example of balsalazide, an anti-inflammatory drug utilised in treating inflammatory bowel disease (IBD), showcases how AGORA2 could be used to inform clinical research, and potentially facilitate personalisation of treatment. Balsalazide has high numbers need to treat (NNT) metrics for inducing remission (NNT:10) and maintenance (NNT:6) in ulcerative colitits^50^, indicating that most patients do not profit from the drug. Consistently, less than half of the analysed microbiomes activated the drug (Figure 5a). Whether this ratio holds in IBD cohorts is yet to be shown, but we revealed that the required azobond reduction is a rare capacity among the gut microbiota, suggesting a role in limiting efficacy of balsalazide treatment. Thus, AGORA2 in conjunction with metagenomics could be utilised to decide on an individual’s benefit of balsalazide treatment. Naturally, follow-up clinical trials would be needed to validate such stratification of inflammatory bowel patients into responders and non-responders. The finding that drug-metabolising capabilities are varying over age-groups, BMI, and sex (Figure 5e, f) demonstrates that AGORA2 in conjunction with community modelling can be utilised in large epidemiological cohort studies, by mapping the drug-metabolising capacities across diseases and risk factors, and thereby opening new research possibilities to understand the role of the microbiome in drug metabolism.

We demonstrated the theoretical drug-metabolising potential of human microbes. Drug response to realistic drug concentrations will require hybrid modelling approaches, e.g., integrating constrained-based modelling with physiological-based pharmacokinetic modelling^51, 52^ and using a constrained-based model of organ-resolved whole-body metabolism with integrated gut microbial community^17^. In a first step, we showed that the diet plays a role for the drug conversion potential of, e.g., sorivudine (Figure S10). Dietary supplements, probiotics, antibiotics, or drugs targeting microbial enzymes, which have been shown to attenuate side effects of drugs^3^, could be predicted and validated using such hybrid modelling approach^51^. Taken together, AGORA2 paves the way for an integrative, multi-scale modelling approach that may enable *in silico* clinical trials^51^ and contribute to precision medicine.

## Author contributions

I.T. and A.H. conceived the study. D.A.R., G.A., and O.E.O. performed comparative genomic analyses. I.T., A.H., S.M., M.H., F.M., J.N.E., and C.S.H. created KBase draft reconstructions. A.H. and S.M. built the semi-automated reconstruction pipeline and the test suite. G.A. and A.H. translated reaction and metabolite identifiers to VMH nomenclature. M.H., G.A., A.H., and F.M. collected experimental data. M.H., F.M., and A.H. collected organism information. M.N. and A.H. formulated the drug module. A.H. performed continuous reconstruction testing and curation. A.H. performed simulations. A.H. and J.H. analysed and visualised the data. J.H. performed statistical analyses. G. P. and R.M.T.F. performed atom-atom mappings. A.H. and J.H. drafted the paper. A.H., I.T., J.H., and D.A.R. edited the paper. I.T. supervised the study.

## Acknowledgements

We thank Prof. Peter Turnbaugh for providing genome sequences for 26 *Eggerthella lenta* strains, Dr. Jan Krumsiek for communicating mouse-associated microbial species, Dr. Cyrille Thinnes for valuable discussions, and Lubin Moussu and Semra Smajic for their help with the comparative genomic effort.

## Material and methods

### Selection of newly reconstructed organisms and retrieval of whole-genome sequences

First, we retrieved 4,185 genomes of human gut-associated strains that were available on PubSEED^53^ (Supplemental Note 6). To expand the species coverage, we performed an extensive literature search of species isolated from or detected in the human microbiome with available whole-genome sequences (Table S1). This search led to the addition of further 1,324 strains, which included 127 genomes of mouse-associated strains. The corresponding whole-genome sequences were retrieved in FASTA format from the NCBI FTP site (ftp://ftp.ncbi.nlm.nih.gov/). Moreover, we included 26 genomes of *Eggerthella lenta* strains^54^ provided through personal communication. Finally, we retrieved 761 human microbial genomes from the Human Gastrointestinal Bacteria Culture Collection (HBC)^44^ in FASTQ format from https://www.ebi.ac.uk/ena/data/view/PRJEB23845 and https://www.ebi.ac.uk/ena/data/view/PRJEB10915. Together with AGORA1.03, which was obtained from the VMH^18^, these combined efforts resulted in 7,206 strains and 1,644 species included in AGORA2.

### Manual refinement of metabolic pathways and gene annotations through comparative genomics

Of the 7,206 analysed strains, 5,438 bacterial strains and three archaeal strains were present in the PubSEED resource^53, 55^ (Supplemental Note 6) and could be re-annotated for their metabolic functions through comparative genomics. 34 metabolic subsystems that had been reconstructed previously for a smaller subset of gut microbial strains^11, 34, 56–58^, as well as a newly created drug metabolism subsystem, were considered for the analysis. All subsystems are available at the PubSEED website.

#### Curation of subsystems

We used the subsystems for biosynthesis of amino acids, B-vitamins, quinones, and nucleotides, as well subsystems for central carbon catabolism, biotransformation of bile acids, respiration, activation of N-acetylglucosamine, fermentation of amino acids, and drug metabolism (Table S2a for a comprehensive list of subsystems). For annotation of the genes in each subsystem, the PubSEED platform was used^53^. Functional roles for each subsystem were annotated based on the (1) prescribed functional role for the protein, (2) sequence similarities of the protein to proteins with previously confirmed functional role, and (3) genomic context (Supplemental Note 7).

#### Metabolic pathways considerations for comparative genomics analysis

Absence of gene(s) for one or more enzymes in a pathway may result in blocked reactions in a metabolic reconstruction. To avoid this, we estimated the completeness of metabolic pathways during the genome annotation. For each potentially synthesised metabolite, all the biosynthetic pathways were collected in agreement with the KEGG PATHWAY resource^59^ and genes of the subsystem were attributed to corresponding steps of the metabolic pathways. Absence of the consequent reactions was determined as a gap. Only pathways with no more than two gaps with gap length of no more than one step (Supplemental Note 8) were further gap-filled and used for generation of reactions.

#### Sequence-based gap-filling

For the gapped pathways, the bidirectional best-hit (BBH) method^60^ was used: (1) The gene corresponding to the gap and present in the genome for the related organisms (belonging to the same species, genus, or family) was used as a query for a BLAST search in the genome with the gap. (2) Possible BBHs were defined as homologs for that alignment with the query protein having an e-value ≤ e-^50^ and protein identity ≥ 50%. (3) For each possible BBH, the reverse search was done for the genome that was a source of the query protein. (4) If the query protein and its best homolog in the analysed genome formed BBH pair, the gap was filled. (5) A similar genomic context for the query protein and its ortholog was considered as an additional confirmation for orthology of the identified BBH pair.

#### Annotation of the drug metabolic genes

To annotate drug-metabolising genes, we used the following pipeline. (1) Identify genes known to encode for drug-metabolising enzymes in a range of microbial organisms, from the scientific literature (Table S5). (2) Using the amino acid sequences of these known drug-metabolising genes as queries, we performed a BLAST search for every analysed genome. (3) The resulting best BLAST hit was then used as a query for the BLAST search in the genome having known drug-metabolising gene to confirm that the known protein sequence and its best BLAST hit form a pair of best bidirectional hits (BBHs). (4) All proteins being BBHs were used for the construction of a rooted maximal-likelihood tree. (5) All previously known proteins were mapped onto the tree, and all monophyletic branches containing known drug-metabolising enzymes were determined (Figure S11). (6) All annotated proteins in these branches were considered as orthologs of the known drug-metabolising proteins. All the proteins not being in branches with known drug-metabolising proteins were considered as proteins with other function and were excluded from further analysis. After the exclusion of the later, a tree was constructed again for orthologs of the known drug-metabolising proteins. (7) For two of the drug-metabolising genes, being the L-tyrosine decarboxylase (TdcA, EC 4.1.1.25) and the cytidine deaminase (cCda, EC 3.5.4.5), we found that genomic context is conserved between species. For these two proteins, we also analysed the genomic context. If genomic context of a candidate gene was similar to that of a known drug-metabolising gene, the candidate was considered as an ortholog of the known protein. Otherwise, it was considered to as a false-positive prediction and excluded from further analysis (Supplemental Note 9, Figure S11). As for (6), the tree was constructed again for only the orthologs of the known proteins. (8) For each tree, including only the orthologs of the known genes, we defined the monophyletic branches containing proteins derived from only one species. For each of such species-specific branch, we predicted subcellular localisation (Supplemental Note 10), using the CELLO v.2.5 system (cello.life.nctu.edu.tw). (9) For cytoplasmic enzymes, drug transporters were predicted based on genomic context (Supplemental Note 11, and Table S2d).

#### Tools

The PubSEED platform^53, 55^ was used to annotate the subsystems. To search for BBHs for previously known proteins, a BLAST algorithm^61^ implemented in the PubSEED platform was used. Additionally, the PubSEED platform was used for analysis of the genomic context. To analyse the protein domain structure, we searched the Conserved Domains Database (CDD)^62^ using the following parameters: an e-value ≤0.01 and a maximum number of hits equal to 500. For the prediction of protein subcellular localisation, the CELLO^63^ web tool was used. Alignments were performed using MUSCLE v.3.8.31^64^. For every multiple alignment, position quality scores were evaluated using Clustal X^65, 66^. Thereafter, all positions with a score of zero were removed from the alignment and the modified alignment was used for construction of the phylogenetic trees. Phylogenetic trees were constructed using the maximum-likelihood method with the default parameters implemented in PhyML-3.0^67^. The obtained trees were midpoint-rooted and visualised using the interactive viewer Dendroscope, version 3.2.10, build 19^68^.

### Literature and database searches

Biochemical and physiological characterisation papers were retrieved by entering the names of AGORA2 species into PubMed (https://www.ncbi.nlm.nih.gov/pubmed/). Information on carbon sources, fermentation pathways, growth requirements, consumed metabolites, and secretion products (Table S3a-e) were subsequently manually extracted on the species and/or genus level from 732 peer-reviewed papers and >8,000 pages of microbial reference textbooks^69^. Moreover, the traits of each reconstructed strain including taxonomy, morphology, metabolism, and genome size were retrieved through database searches. The taxonomic classification of the strains was retrieved from NCBI Taxonomy (https://www.ncbi.nlm.nih.gov/taxonomy/) and, to our knowledge, is up to date at the time of this publication. Information on morphology, habitat, body site, gram status, oxygen status, metabolism, motility, and genome size were retrieved from the Integrated Microbial Genomes and Microbiomes^70^ database (https://img.jgi.doe.gov/) (Table S1).

### Generation of draft reconstructions

Draft reconstructions were generated through the KBase^12^ narrative interface. Genomes present in KBase were directly imported into the narrative. Otherwise, genomes in FASTA format were uploaded into the Staging Area and subsequently, imported into the narrative through the “Batch Import Assembly From Staging Area” (narrative.kbase.us/#catalog/apps/kb_uploadmethods/batch_import_assembly_from_staging) app. Genomes in FASTQ format were directly imported into the narrative through the “Import Paired-End Reads From Web” (https://narrative.kbase.us/#catalog/apps/kb_uploadmethods/load_paired_end_reads_from_URL) app after retrieving the links to the corresponding files from https://www.ebi.ac.uk/ena/data/view/PRJEB23845 and https://www.ebi.ac.uk/ena/data/view/PRJEB10915. The imported assemblies were annotated using RAST subsystems^71^ through the “Annotate Multiple Assemblies” (https://narrative.kbase.us/#appcatalog/app/RAST_SDK/annotate_contigsets) app. Draft metabolic reconstructions were generated through the “Create Multiple Metabolic Models” (https://narrative.kbase.us/#appcatalog/app/fba_tools/build_multiple_metabolic_models) app and exported in SBML format through the “Bulk Download Modelling Objects” (https://narrative.kbase.us/#appcatalog/app/fba_tools/bulk_download_modeling_objects) app.

### Semi-automated, data-driven refinement pipeline

The AGORA pipeline has been described previously^11^. Here, we revised the pipeline substantially to accommodate additional curation efforts needed for the new reconstructions. Specifically, we (i) translated ~1,000 additional reactions and ~800 metabolites from KBase to VMH^18^ nomenclature; (ii) introduced additional gap-filling reactions, where needed, to enable biomass production under anoxic conditions on the previously defined Western diet^11^ with thermodynamically consistent reaction directionalities; (iii) removed futile cycles resulting in thermodynamically implausible ATP production by making the responsible reactions irreversible; (iv) ensured through gap-filling and/or deletion of appropriate reactions that all reconstructions captured the collected experimental data (Table S3a-c); and (v) adjusted biomass objective functions to account for class-specific cell membrane and cell wall structures (Supplemental Note 3). As described previously^11^, all solutions were manually determined for few reconstructions and subsequently propagated to many reconstructions, as appropriate. Reactions identified through comparative genomics (Table S2b-c) were added to the up to 5,438 reconstructions. Non-gene associated reactions, for which the respective gene could not be found through comparative genomics, were removed from the draft reconstructions if doing so did not abolish biomass production. Moreover, published information on metabolite uptake and secretion in ~570 gut microbial species retrieved from^72^ was mapped onto VMH nomenclature and used for validation of the predictive potential and subsequent further expansion of the reconstructions (Supplemental Note 2). Finally, to ensure proper compartmentalisation, a periplasm compartment was introduced. For consistency, the existing 818 AGORA1.03 reconstructions (version 25.02.2019, available at https://www.vmh.life/files/reconstructions/AGORA/1.03/AGORA-1.03.zip) also underwent the revised pipeline. The AGORA1.03 reconstruction of *Staphylococcus intermedius* ATCC 27335 was removed since it was a duplicate of the newly reconstructed strain *Streptococcus intermedius* ATCC 27335. The names of 26 AGORA 1.03 reconstructions were changed to account for recent changes in taxonomical classification (Table S1).

All pipeline functions were written in MATLAB (Mathworks, Inc.) version R2018b and relied on functions implemented in the COBRA Toolbox^36^. All newly included metabolites and reactions were formulated based on literature and/or database^18, 73, 74^ searches while ensuring mass and charge balance through the reconstruction tool rBioNet^75^.

### Test suite for quality control and quality assurance

To ensure that curation efforts were successful, a COBRA Toolbox-based test suite for the AGORA2 reconstructions was created and regularly performed during revision of the pipeline. Specifically, it systematically accessed whether each reconstruction (i) grew anaerobically on the Western diet, (ii) had correct reconstruction structure, i.e., mass and charge balance, and correct syntax for gene-protein-reaction associations, (iii) was thermodynamically feasible, e.g., produced realistic amounts of ATP, and (iv) captured known metabolic traits of the organism according to the collected experimental and comparative genomic data. Table S4 summarises all features that were tested by the test suite.

The comparison with the experimental and comparative genomic data was carried out by predicting the capability of AGORA2 reconstructions to take up or produce reported consumed or secreted metabolites. If the strain was known to take up or produce the metabolite and the corresponding AGORA2 reconstruction could also take or secrete the metabolite, this resulted in a true positive prediction, while a false negative prediction occurred when the strain was known to have this capability but the corresponding reconstruction did not capture the trait. For growth requirements, two types of experimental information were available: nutrients that are known to be required by the organism in question, and nutrients that are known to not be required. This information allowed us to additionally determine true negatives (the nutrient is nonessential for growth in the experiment and in the reconstruction) and false positives (the nutrient is nonessential for growth in the experiment but required for growth in the reconstruction) for growth requirements. False positive and false negative predictions were routinely retrieved, and reaction gap-filling and/or deletion solutions were included in the reconstruction pipeline functions to eliminate them. This refinement was performed in an iterative effort.

### Flux and stoichiometrically consistent reactions

The subset of flux and stoichiometrically consistent reactions, as defined in^20^, was retrieved for each AGORA2 reconstruction and corresponding draft reconstruction through the ‘findFluxConsistentSubset’ and ‘findStoichConsistentSubset’ functions implemented in the COBRA Toolbox^36^. The fraction of stoichiometrically and flux consistent reactions, excluding exchange and demand reactions, was determined for each draft and curated reconstruction. Briefly, the subset of stoichiometrically consistent reactions in a reconstruction includes all reactions that are mass and charge conserved, excluding exchange reactions, which are by definition mass and charge imbalanced^20^. The subset of flux consistent reactions consists of all reactions that are stoichiometrically consistent and can carry flux^20^.

### Formulation of the drug reactions

A literature search for microbial enzymes known to transform, degrade, activate, inactive, or indirectly influence commonly prescribed drugs was performed yielding 15 enzymes in total (Figure 3a, Table S5), which are encoded by 29 genes (Table S2b). To enable comparative genomic analyses, only drug transformations that could be linked to specific protein-encoding genes were considered. As described above, enzyme-encoding genes were analysed in their genomic context as outlined in^76^ using PubSEED subsystems^19, 53^.

Literature and database searches were performed for the metabolic fate of commonly prescribed human-targeted drugs. The structures of 287 drug metabolites and drug degradation products were retrieved from 73 peer-reviewed papers, HMDB^77^, DrugBank^78^, and Transformer^79^. Reactions were formulated based on the collected experimentally determined drug structures, drug downstream product metabolite structures, and reaction mechanisms. Both, cytosolic and extracellular, enzymatic reactions were formulated depending on the identified subcellular protein locations. Since at least six drugs undergoing glucuronidation in the human body have been shown to be substrates for the microbial ß-glucuronidase^80, 81^ (Table S5), it was assumed that all retrieved glucuronidated drug metabolites (118 in total) could serve as substrates. Additionally, ß-glucuronidase reactions were formulated for 33 glucuronidated drug metabolites from a previously reconstructed module of human drug metabolism^82^. New metabolites and reactions were assigned VMH IDs following standards in nomenclature used for COBRA reconstructions^8^, and formulated while ensuring mass and charge balance through the reconstruction tool rBioNet^75^. In total, for 98 drugs (Figure 3b), 353 unique metabolites, 381 enzymatic reactions, 373 exchange reactions, and 710 transport reactions (Table S6a-b) were formulated.

### Atom-atom mapping

Atom-atom mappings were obtained using a database standardisation pipeline described in ^83^ and the AGORA2 reconstructions, specifically the information of the metabolites present in the reconstructions together with the reaction stoichiometry. The database standardisation pipeline^83^ was executed in MATLAB and use different external software tools, such as ChemAxon^84^, Open Babel^85^ and, the Reaction Decoder Tool^86^. The process to obtain the atom-atom mappings for the AGORA2 reconstructions can be summarised as follows: 1) 1,838/3,533 metabolic structures of the metabolites present in the AGORA2 reconstructions were collected from different chemical databases, such as VMH^18^, KEGG^74^, HMDB^77^, PubChem^87^ and ChEBI^88^ databases. The metabolic structures were standardised based on the InChI algorithm^89^ and can be found in the VMH database^86^; 2) the standardised metabolites and the reaction stoichiometry in the AGORA2 reconstructions were used to generate 5,583/7,300 MDL RXN files; 3) 5,583/7,300 AGORA2 reactions were atom mapped using the Reaction Decoder Tool algorithm^86^ for active reactions and a pipeline’s algorithm^83^ for passive transport reactions and coupled transport reactions. Atom-atom mappings can be found in the VMH database^18^.

### Simulations

All simulations were performed in MATLAB (Mathworks, Inc.) version R2018b with IBM CPLEX (IBM) as the linear and quadratic programming solver. The simulations relied on functions implemented in the COBRA Toolbox^36^, and the Microbiome Modelling Toolbox^37^. Flux balance analysis (FBA)^26^ was used to interrogate drug metabolism. All additional scripts for data generation, data analysis, and data visualisation are available at https://github.com/ThieleLab/CodeBase.

### Validation of drug-metabolising capacities against independent, experimental data

A literature search was performed for *in vitro* experiments demonstrating the capabilities of human microbial strains to metabolise reconstructed drugs through the 15 annotated enzymes (Table S7). If no studies on the reconstructed drugs were found for the enzyme, studies on demonstrated drug activity of the enzyme were recorded. Subsequently, the capabilities to metabolise the drugs through the respective enzymes for 169 AGORA2 metabolic model, for which data could be found, were tested by computing whether the corresponding reaction could carry flux (Table S7). If possible, the tested organisms were matched to AGORA2 models on the strain level, otherwise pan-species models were used. Accuracy, sensitivity, and specificity of predictions were calculated by determining the number of true positive, true negative, false positive, and false negative predictions.

### Drug yields

To determine each strains’ capability to metabolise drugs, all AGORA2 were constrained with a simulated Western diet^11^ and the flux through the exchange reactions corresponding to each drug was minimised through FBA, corresponding to maximal uptake of the drug. For all AGORA2 organisms capable to take up at least one drug, the yield of ATP, carbon, and ammonia from 1 mmol of the drug/g_dry weight_/hr was evaluated as follows. Each reconstruction was constrained to only allow the uptake of water, phosphate, and oxygen (VMH IDs: h2o, pi, o2). Demand reactions for ammonia as well as CO_2_ and pyruvate (as proxies for carbon sources) (VMH IDs: nh4, CO_2_, pyr) were added, while a demand reaction for ATP (VMH ID: atp) already existed in each reconstruction. Next, the uptake of each drug metabolite (15 in total, one representative for each enzyme) was allowed one by one at an uptake rate of 1 mmol/g_dry weight_/hr. For each drug metabolite, the yields of ATP, ammonia, CO_2_, and pyruvate from each drug metabolite were computed using flux balance analysis (FBA) by maximising the flux through the respective demand reactions. As control, yields were also computed for 1 mmol/g_dry weight_/hr of glucose and without any metabolites added.

### Pairwise simulations

We randomly selected 50 AGORA2 strains each out of (i) all 872 strains that could use glucuronic acid but did not have the b-glucuronidase enzyme, (ii) all 79 strains that did have b-glucuronidase but could not use glucuronic acid, (iii) all 1,473 strains that could use glucuronic acid and had b-glucuronidase, and (iv) all 3,423 strains that neither used glucuronic acid nor had ß-glucuronidase. The resulting 200 strains were joined in all possible combinations, as described previously^90^ resulting in 19,9000 pairwise models. For each pair, co-growth was predicted using functions implemented in the pairwise modelling module in Microbiome Modeling Toolbox^37^ with the possible outcomes being competition, parasitism, amensalism, neutralism, commensalism, and mutualism as defined in^90^. Pairwise models were grown anaerobically on the previously defined Western diet^11^ under three conditions: (i) no additional compound added, (ii) supplementation with 10 mmol/gdry weight/hr of irinotecan in its glucuronidated form SN38G (VHM ID: sn38g), (iii) supplementation with 10 mmol/gdry weight/hr of free glucuronic acid (VMH ID: glcur).

### Definition of an average Japanese diet

An average Japanese diet was defined based on the mean daily food consumption in 106 Japanese dietitians determined from a food frequency questionnaire and 28 days weighed diet records^91^. The reported food items were mapped to the corresponding or closest possible item in a database of >8,000 food items available on the VMH^18^ website. The Diet Designer function on the VMH website permits the design of a personalised diet through input of daily food consumption quantities with the uptake flux values in mmol/person/day for each nutrient as the output^18^. The designed diets are suitable for the contextualisation of personalised microbiome models^37^. The mapped mean daily food consumption quantities in gram were entered into the Diet Designer tool and the generated diet uptake fluxes were exported. To perform microbiome modelling simulations, the retrieved diet fluxes were adapted to enable biomass production of all AGORA2 strains by using the ‘adaptVMHDiet2AGORA’ function as described previously^34^. Table S13a) shows the retrieved quantities of food items in grams that were the input for the Diet Designer tool. Table S13b shows the adapted uptake fluxes in mmol/person/day that were used to contextualise the microbiome models for 616 Japanese individuals *in silico* (see also next section).

### Simulation of drug metabolism by individual gut microbiomes

Previously, metagenomic sequencing from faecal samples of a cohort of 616 Japanese colorectal cancer patients and healthy controls had been performed^28^. Species-level abundances for this cohort, which has been determined with MetaPhIAn2^92^, were retrieved from https://www.nature.com/articles/s41591-019-0458-7#MOESM3. Unclassified taxa on the species level, eukaryotes, and viruses were excluded. Of the remaining 517 species, 502 (97%) could be mapped onto 1,644 AGORA2 species based on names. Pan-species models for AGORA2 were created through the ‘createPanModels’ function. From the pan-species models, personalised microbiome models for each of the 616 samples were built and parameterised as described elsewhere^34, 37^ with the species-level abundances as input data. To contextualise the models with appropriate diet constraints, the Average Japanese Diet described above was used instead of the previously used Average European diet^34^. To predict the drug conversion potential of each microbiome, the faecal secretion reactions for 13 drug metabolism end products were optimised one by one using FBA^26^, while providing the respective precursor drug as well as oxygen at a de facto unlimited uptake rate of 1000 mmol/g_dry weight_/hr.

### Shadow price analysis

To determine species in microbiome models that were of importance for the microbiome’s combined potential to metabolise a drug, a shadow price analysis was performed as described previously^34^. Briefly, shadow prices are a feature of every flux balance analysis solution (i.e., the shadow price is the dual to the primal linear programming problem) that reflect the contribution of each metabolite in the model to the flux through the objective function^7^. Briefly, a non-zero shadow price for a metabolite indicates that this metabolite has importance for the total flux capacity through the optimised objective function, i.e., in our case, the secretion of a drug metabolic product. A shadow price of zero indicates that increasing the availability of this metabolite would not change the flux through the objective function. To determine the species that were bottlenecks for the conversion potential of the 13 drugs in each microbiome model, nonzero shadow prices for species biomass metabolites (‘species_biomass[c]’), which reflect the contribution of the species to the community biomass reaction, were retrieved.

### Statistical analysis

We analysed statistically the net production capacity of 13 drug metabolites (Figure 5b) among 252 healthy individuals and 364 CRC patients. For each drug metabolite, we calculated the mean flux and the share of microbiomes with a flux greater zero. Drug metabolites, which had in over 50% of the cases a zero flux were dichotomised (can be produced vs. cannot be produced) and subsequently, analysed via logistic regressions. Drug metabolites with over 50% non-zero entries were analysed via linear regressions using heteroscedastic robust standard errors. First, we investigated potential effects of basic covariates (age, sex, and BMI) via generalised linear regressions (logistic or linear) with the net production capacity being the response variable (dichotomised or metric). Age and BMI were introduced into the models as restricted cubic splines^93^ using four knots (the 5%-percentile, the 33%-percentile, the 66%-percentile, and the 95%-percentile) resulting in three spline variables, each to test on potential non-linear relationships. Significance was then determined by testing the three spline variables belonging to age (or BMI, respectively) simultaneously on zero via the Wald test^93^. While for age substantial non-linearities were found, no indication for non-linear BMI effects could be identified. The final models included, therefore, only the linear BMI term. Second, we tested for potential associations of net production capacities with case control status. This was done via generalised linear regressions (logistic or linear) with the net production capacity being the response variable (dichotomised or metric), while adjusting for age (restricted cubic splines), sex (male/female), and BMI (linear). Finally, we tested analogously on associations with the CRC stage by introducing the stage as categorical variable (multiple polyps, 0, I/II, and III/IV) into the model. We corrected for multiple testing using the false discovery rate, adjusting significance values for 13 tests per analyses stream. A test was considered nominal significant with p<0.05 and FDR-corrected significant if FDR<0.05. For sensitivity analysis, we recomputed the drug-metabolising capabilities using an average European diet instead of a Japanese diet. Then, we calculated Pearsons correlations for each drug metabolite between the secretion potentials under Japanese and an average European diet. All statistical analyses were performed with STATA 16/MP.

### Data visualisation

The phylogenetic tree of AGORA2 organisms was constructed in PhyloT (https://phylot.biobyte.de/) and visualised in iTOL (https://itol.embl.de/)^94^. Violin plots were generated in BoxPlotR (http://shiny.chemgrid.org/boxplotr/). Clustering of taxa by reaction presence through t-distributed stochastic neighbour embedding (t-SNE)^95^ was performed using the t-SNE implementation in MATLAB with Euclidean distance, barneshut set as the algorithm, and perplexity set to 30. Taxa with fewer representatives than 0.5% of all clustered strains were excluded from the t-SNE plots. Circle plots were generated using the online implementation of Circos^96^. Figure 5 was generated with the graphics functions of STATA 16/MP. All other data was visualised in MATLAB and R^97^.

