## Supplemental Material for "AGORA2: Large scale reconstruction of the microbiome highlights wide-spread drug-metabolising capacities"

**Supplemental Note 1: AGORA 2.0 represents a substantial expansion in size and scope over AGORA 1.03**

We aimed to extend the scope of AGORA2 beyond cultured and gut-associated strains found in Western microbiomes. While AGORA 1.03 mostly consisted of cultured species, AGORA2 includes 487 currently uncultured and/or uncharacterised strains (Figure 3a, Table S1). For instance, AGORA2 contains 737 bacterial isolates, including 105 novel species, from the Human Gastrointestinal Bacteria Culture Collection (HBC)[1]. AGORA2 also accounts for body sites other than gut, such as skin and mouth (Table S1). While the selection of AGORA 1.03 strains was mainly based on species found in healthy young Western individuals[2], AGORA2 also accounts for species detected in a cohort of elderly Parkinson's Disease patients and controls[3], a cohort of Japanese colorectal cancer patients and controls[4] as well as species isolated from Polynesian, Saudi, and Senegalese individuals[5]. Finally, we expanded the coverage of AGORA beyond human-associated strains by reconstructing 127 mouse-associated strains (Figure 3a, Table S1), thus, enabling modelling of the mouse microbiome. Mouse models are an important tool in microbiome research, but the microbiome is distinct from human[6]. It should be noted that Bacilli and Gammaproteobacteria were overrepresented in the strain selection (Figure 3b) reflecting a likely sequencing bias for well-studied species and/or opportunistic pathogens.

**Supplemental Note 2: The data-driven reconstruction pipeline results in high-quality reconstructions that capture known biochemical and physiological properties**

The AGORA2 reconstructions underwent continuous testing through a test suite that ensured correct reconstruction structure, biochemical and thermodynamic consistency, as well as good agreement with experimental and genomic findings and known traits of the organism (Table S4). These efforts ensured that all curated reconstruction-derived condition-specific metabolic models could grow anaerobically on the previously defined Western diet[7] and produced realistic amounts of ATP, which was not the case for models derived from most draft reconstructions (Figure 3c). We collected information on carbon sources, fermentation pathways, and growth requirements (Table S3a-c) that served as input data for the pipeline. Using an iterative approach, the semi-automatically curated reconstructions were continuously tested against the input data (Figure 1). Discrepancies between experimental data and model predictions identified by the test suite were manually inspected and corrected (Methods). As a result, while the metabolic models derived from draft reconstructions showed low prediction accuracy, those derived from the curated AGORA2 reconstructions, as expected, agreed very well with the experimental data (Figure 2b). For instance, defined media had recently been reported for 74 AGORA2 strains[8] and all 74 reconstructions could grow on the respective media.

The metabolic reconstructions were further curated by mapping a published compendium of metabolite uptake and secretion data for ~570 human microbes[9] onto the AGORA2 strains (Table S3d-e). Corresponding exchange and transport reactions were added for each microbe reported to take up and/or secrete a metabolite. At this point, the metabolite uptake and secretion capabilities of models derived from the AGORA2 semi-automatically curated reconstructions were tested against the uptake and secretion data compendium and good agreement was found (94% and 86% of all reconstructions-derived models agreeing with metabolite uptake and secretion data, respectively). Thus, due to the extensive data-driven curation and refinement already performed, AGORA2 captured species-specific catabolic and biosynthetic pathways present in the human gut microbiome very well. Afterwards, the reconstructions were further improved by performing gap-filling to ensure the uptake and secretion of metabolites reported in[9], resulting in an final agreement with metabolite uptake and secretion data of 99% for both.

**Supplemental Note 3: Curation of biomass objective functions**

Gram-positive and -negative bacteria differ in their cell wall structure[10]. Gram-positive bacteria possess a thick layer of teichoic acid and an inner but no outer membrane, while gram-negative bacteria have an inner and an outer membrane separated by a large periplasmatic space with the outer membrane carrying lipopolysaccharides (LPS)[10]. As exceptions, the Deinococcus-Thermus phylum has on outer membrane, but does not have LPS[11], certain Firmicutes, such as *Acidaminobacter* and *Gracilibacter* sp. stain gram-negative, but have a gram-positive cell wall structure[10, 12], and the Chloroflexi phylum stains gram-negative, but has no LPS and no outer membrane[10]. Moreover, the Tenericutes phylum does not possess a cell wall[10] and archaea have ether lipids in their membranes instead of the bacterial ester lipids[10]. When inspecting the biomass objective functions (BOFs) in the draft reconstructions, it was found that 32% of reconstructions of gram-positive organisms had gram-negative components in their BOF or vice versa. Moreover, 35 Tenericutes and seven Archaea draft reconstructions incorrectly had standard bacterial cell wall components in their BOFs. To correct this, all BOFs were checked based on the respective organisms' gram status and corrected by removing incorrect metabolites and add teichoic acid or LPS, as appropriate. For gram-negative organisms with an outer membrane, a periplasmatic compartment was added. The exceptions listed above were taken into account. An archaeal BOF was formulated by retrieved ether lipid structures and the corresponding biosynthesis reactions from the reconstruction of *Methanosarcina barkeri*[13]. Gap-filling reactions enabling cell wall component production were also added to the corresponding reconstructions, if necessary.

**Supplemental Note 4: Reconstruction features across taxa**

The reaction content of the 23 classes with the highest numbers of representatives in AGORA2 was visualised through t-distributed stochastic neighbour embedding (t-SNE)[14] (Methods, Figure 3e-g). Generally, reconstructions clustered by class indicating that related organisms were similar in reconstruction content (Figure 3e). However, multiple subclusters of microbial classes were observed, especially for the Bacilli and Gammaproteobacteria classes (Figure 3e). Clustering Bacilli and Gammaproteobacteria representatives separately revealed multiple subclusters in the *Enterococus*, *Staphylococcus*, and *Streptococcus* genera (Figure 3f) as well as in the *Escherichia* and *Salmonella* genera (Figure 3g). These clusters were already observed in the KBase draft reconstructions (Figures S9a-c) indicating that this was a result of the genome annotation.

The number of reactions, metabolites, and genes per reconstructions also varied by taxon (Figure 3h-j). The highest numbers of reactions, metabolites, and genes were found in the Proteobacteria phylum, likely as a result of many representatives of this phylum having large genomes (Table S1) and a generalist type of metabolism[10]. The smallest reconstructions were found in the Tenericutes phylum, which consisted of organisms without a cell wall[10]. Growth rates on the anaerobic Western diet also correlated with reconstruction size and were in a biologically realistic range (Figure 3k).

### Supplemental Note 5: Interspecies interactions and bottlenecks in microbial drug metabolism

For some drug metabolic products, only a subset of microbiome samples correlated with the abundances of the enzymes directly producing them (Figure S6). We hypothesised that this was due to multiple species being involved in the metabolism of these drugs. An interspecies interaction in the metabolism of the Parkinson's Disease drug levodopa has been demonstrated previously[15]. To identify species and enzymes that were bottlenecks for drug metabolism, we performed a shadow price analysis as described previously[16] (Methods). Non-zero shadow prices of biomass metabolites indicated that these species were flux bottlenecks for the drug metabolic product (Methods, Figure 6, Table S9). For instance, for the case of balsalazide conversion into its active form by azoreductase, and subsequent conversion into N-acetyl-5-aminosalicylic acid by arylamine-N-acetyltransferase, the abundance of azoreductase-carrying species was always the rate-limiting step (Figure 6a). Only seven species including *Clostridium perfringens*, *Gordonibacter pamelae*, and *Pseudomonas aeruginosa* were relevant for balsalazide conversion flux in any microbiome demonstrating the rarity of this function (Figure 6a, Table S9). Taken together, mechanistic modelling revealed in a microbiome-specific manner the species serving as bottlenecks for desirable or undesirable drug conversions. Similarly, for the conversion of the anticancer prodrug 5-fluorocytosine to the active drug 5-fluorouracil via cytosine deaminase, and the detoxified end product 5,6-dehydro-5-fluorouracil via dihydrouracil dehydrogenase, either the first or the second step were flux bottlenecks (Figure 6b). Examples are *Anaerotruncus colihominis* and *Hungatella hathewayi*, which carry only cytosine deaminase, and *Escherichia coli*, which has both cytosine deaminase and dihydrouracil dehydrogenase (Figure 6b). Finally, for the previously described case of levodopa metabolism[15], two variations of flux limitations could be distinguished. If production of the end product m-tyramine correlated directly with dopamine dehydroxylase abundance, the abundance of *Eggerthella lenta*, the only species carrying this enzyme was flux-limiting (Figure 6c). In contrast, for microbiomes outside the correlation curve, the abundance of species carrying tyrosine decarboxylase, which produces dopamine from levodopa (e.g., *Enterococcus* sp.), was flux limiting (Figure 6c). A species-species interaction in levodopa metabolism involving *Enteroccocus* sp. and *Eggerthella lenta* had been previously reported[15]. Since this microbial pathway lowers the bioavailability of the active drug levodopa[15], identifying the species that serve as bottlenecks for levodopa conversion is an important consideration for Parkinson disease treatment strategies.

### Supplemental Note 6: Retrieval of genomes for human gut-associated strains

An initial set of genomes for a comparative genome analysis included 632 of 773 genomes from the AGORA genome set[7]. To extend this set, we retrieved genomes for all the strains for microbial species associated with human gut[17], and the genomes for 4,881 of such strains were available at the PubSEED resource. To check the quality of genome sequencing and assembly, we analysed the distribution of 31 genetic marker genes that are these proteins because they are nearly universally distributed in Bacteria and exist as single copy genes within each genome[18]. In agreement with the distribution of the genetic marker genes, this gene list was modified. Thus, the genes *dnaG*, *infC*, and *pgk* were excluded because these genes often exist as multiple non-identical copies within certain genome, whereas the gene *pyrG* was excluded because its absence in multiple analysed genomes. Based on the distribution of the other genetic marker genes, 43 genomes were excluded out of the genome set as lacking multiple genetic marker genes and/or the presence of multiple identical copies of the gene in a single genome. Thus, genome set used for the comparative genomics manual refinement, included 4,848 genomes. For an analysis of drug metabolism, the genome set was even more extended by addition of 643 genomes for other gut associated strains that had been retrieved through literature searches (Supplemental Note 1) and were available in the PubSEED database. Thus, for drug metabolism, genome set was extended up to 5,438 microbial genomes.

**Supplemental Note 7: Curation of the subsystems, annotation of protein functional roles**

All the functional roles corresponded to a single catalysed reaction, or a set of catalysed reactions (see below), were grouped into subsets, if possibly. For example, NADH- and NADPH- specific forms of FMN-dependent azoreductases (EC 1.7.1.6 and EC 1.7.-.-, respectively) were attributed to different subsets. Two types of the subsets were created, (1) including all subunits for certain enzyme or transporter, such as catalytic and electron-transfer subunits of digoxin reductase, and (2) including alternative names for the same enzymatic activity, such as b-galactosidases (EC 3.2.1.23) of the families GH2, GH35, and GH42. The single protein can belong to more than one subset, for example, NAD(P)H-specific form of azoreductase was attributed to subsets for both NADH- and NADPH-specific forms, nonetheless this protein is not homologous to any of two previous.

To include all the alternative names for the same enzyme or transporter, chromosomal gene clusters were analysed. This was done for all genes that were absent in a certain genome but were present genomes of organisms belonging to the same species or genus. The following procedure was applied: (1) Genome context of the analysed gene was compared and genes, clustered with the analysed gene in multiple genomes of the related organisms were determined. (2) Orthologs of the clustered genes were searched in the analysed genome, and, if they were found, (3) their genomic context in the analysed genome was studied to find a gene that (i) have a genome context similar to that was observed in the genomes of the related organisms for the analysed gene and (ii) have a name synonymous to the name of the analysed gene or other genes in the subset including the analysed gene. (4) If such candidate was found, a new functional role was included into the subsystem and included into the corresponding subset.

**Supplemental Note 8: Estimation of completeness of the analysed metabolic pathways**

Most of the curated subsystems correspond to biosynthesis of certain metabolite(s). All the biosynthetic pathways for each of potentially synthesised metabolite were collected and analysed. More than one pathway may be possible for a single metabolite, as well the same gene can be included into different pathways for the synthesis of different metabolites.

Based on the presence of enzymatic genes, all the pathways were classified to the following categories. (1) Complete pathways included genes for all the enzymes of the pathway. (2)

Gapped pathways were defined as that have no genes for no more than two reactions and length of each gap in the pathway does not exceed one reaction. (3) For incomplete pathways genes for more than two reactions were absent or extension of a gap exceeded two or more reactions. (4) If no gene of a pathway was present in the genome, the pathway was defined as absent. For the incomplete pathways, no reactions corresponding to the present genes were included into metabolic reconstructions. For the gapped pathways, a gap-filing procedure was applied, whereas for the complete pathways, reactions for all the genes were included into the reconstructions. The metabolite was considered as being synthesised by a microorganism if at least one complete or gapped biosynthetic pathway was predicted. This classification was applied to all analysed pathways except the (1) drug and bile acids transformations, which have been shown to rely on microbial collaboration to complete a pathway[15]; (2) respiration because all the curated pathways consist of only one reaction, and (3) central carbon catabolism because these pathways are tightly connected to the biosynthetic pathways and may be defined as incomplete whereas not generating blocked reactions[7].

**Supplemental Note 9: Manual refinement of the annotations for the drug-metabolising enzymes**

For a refinement of functional annotations for the drug-metabolising enzymes the following procedure was applied. (1) Phylogenetic tree was constructed for protein sequences of all the found BBHs. (2) Experimentally confirmed drug-metabolising enzymes were mapped on the tree. (3) Monophyletic branches containing the experimentally confirmed enzymes were defined. (4) Branches lacing these enzymes were considered as false-positive predictions and were excluded out of the analysis (Figure S9). Additionally, it was found that two of the drug-metabolising enzymes have a conserved genomic context. Thus, a gene for L-tyrosine decarboxylase (TdcA, EC 4.1.1.25) is clustered together with a gene for Tyrosyl-tRNA synthetase (EC 6.1.1.1) whereas a gene for Cytidine deaminase (cCda, EC 3.5.4.5) is clustered together with genes for Pyrimidine-nucleoside phosphorylase (EC 2.4.2.2) and Deoxyribose-phosphate aldolase (EC 4.1.2.4). Such a genomic context was used to distinguish between genes for the drug-metabolising enzymes (clustering is conserved) and their paralogs (no clustering).

**Supplemental Note 10: Analysis of subcellular localisation of drug-metabolising enzymes**

For all the drug-metabolising enzymes the following procedure was applied. (1) For all the enzymes homologous to each other, maximal-likelihood phylogenetic tree was constructed; for example, for β-galactosidases three trees were constructed, for the GH2, GH35, and GH42 families, respectively. (2) For every tree, species-specific monophyletic branches were defined. (3) Subcellular localisation was predicted with CELLO[19, 20] web tool for one randomly selected protein for every species-specific monophyletic branch and then extrapolated to the whole species-specific monophyletic branch.

**Supplemental Note 11: Prediction of drug transporting proteins**

Cytoplasmic drug metabolising enzymes require the presence of transporters delivering corresponding drugs into the cytoplasm. To predict these transporters, we analysed genomic context for the predicted cytoplasmic enzymes. Candidate drug transporters should satisfy the following criteria: (1) Genes for these transporters should be chromosomally co-localised with the genes for the cytoplasmic enzymes. (2) This co-location should be evolutionary conserved, i.e., should be observed in more than one species. (3) Genes for candidate drug transporters should demonstrate sequence similarity with known domains specific for transport proteins, which was checked by search on CDD database[21] using the following cut off parameters: an e-value ≤0.01 and a maximum number of hits equal to 500.

### Supplemental Figures

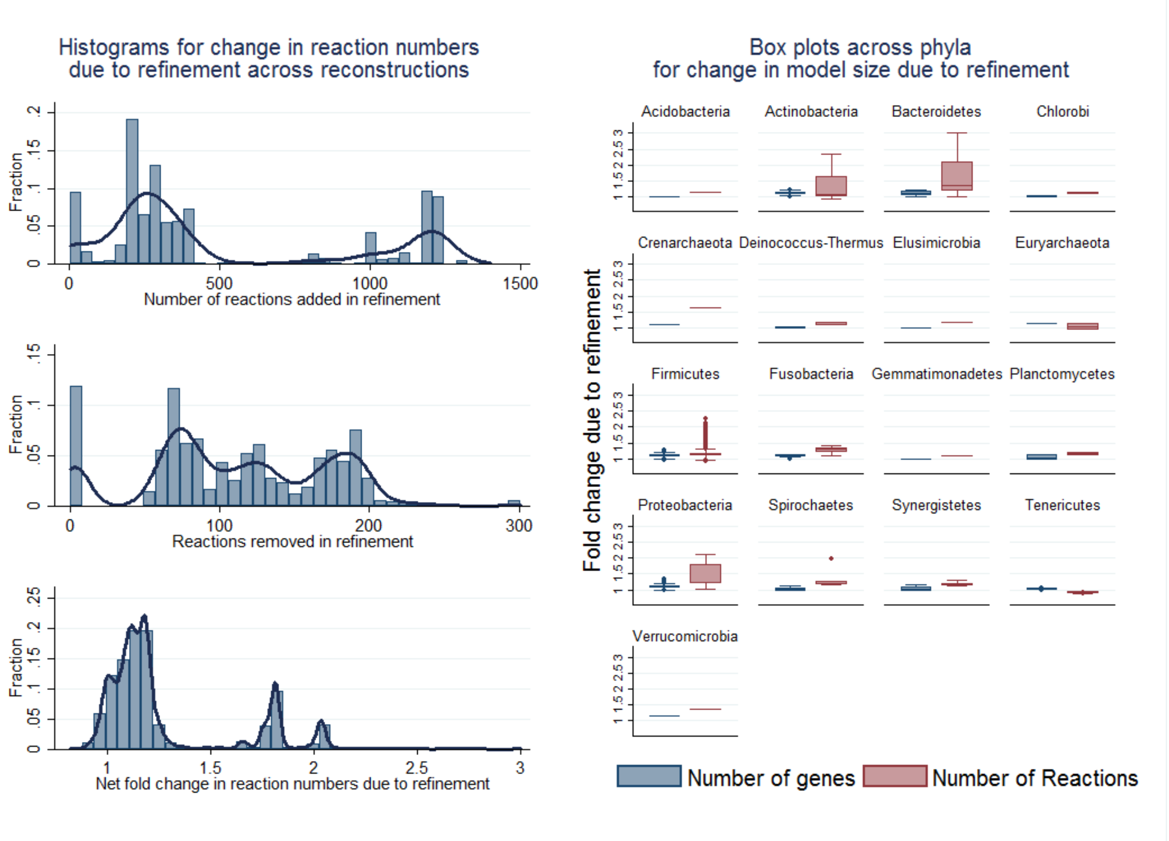


**Figure S1:** Change in model size due to refinement of reconstruction for the 5,438 reconstructions, for which comparative genomics were performed. **Left Panel**: Histograms for change in reaction numbers across reconstructions. **Right panel**: Box plots across phyla for fold change in gene and reaction numbers due to reconstruction. High numbers of added reactions were result of adding drug reactions to reconstructions with drug-metabolising potential.

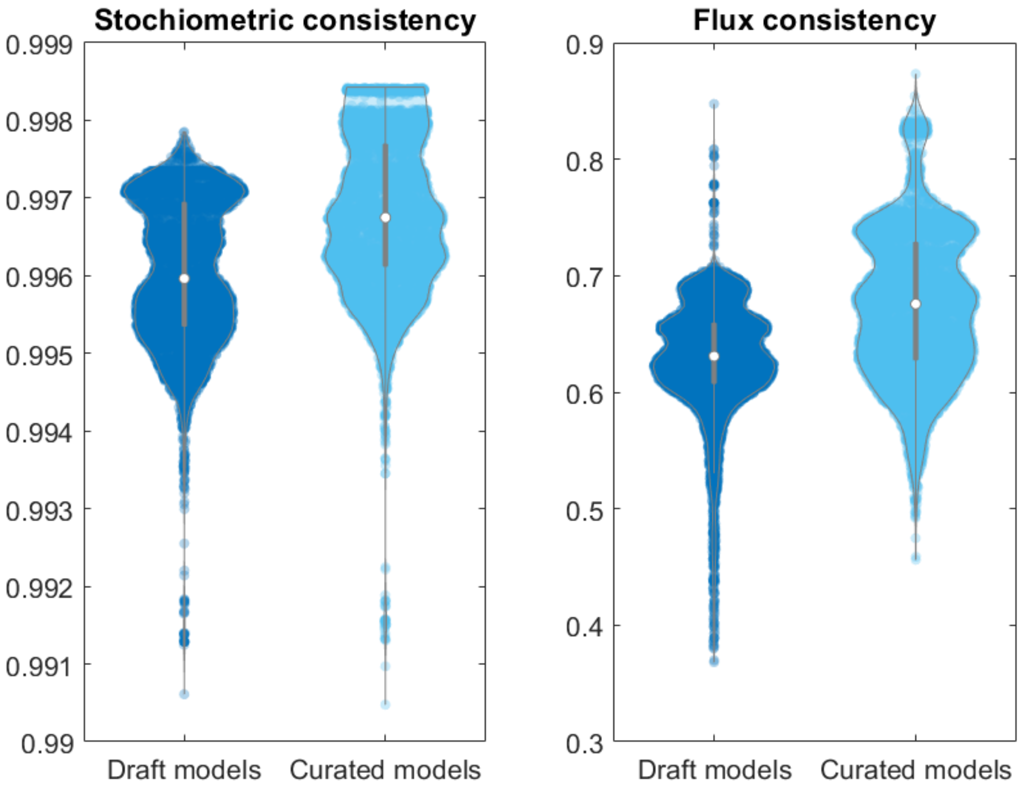


**Figure S2**: Fraction of reactions that are stoichiometrically and flux consistent as defined in [22] for the 7,206 AGORA2 draft and curated reconstructions (mean = 67.6%, 95%-CI(67.4%,67.8%) for AGORA2, mean= 62.9%, 95%-CI(62.8%,63.1%) for the draft reconstructions). Exchange and demand reactions, which are stoichiometrically inconsistent by definition, were excluded.

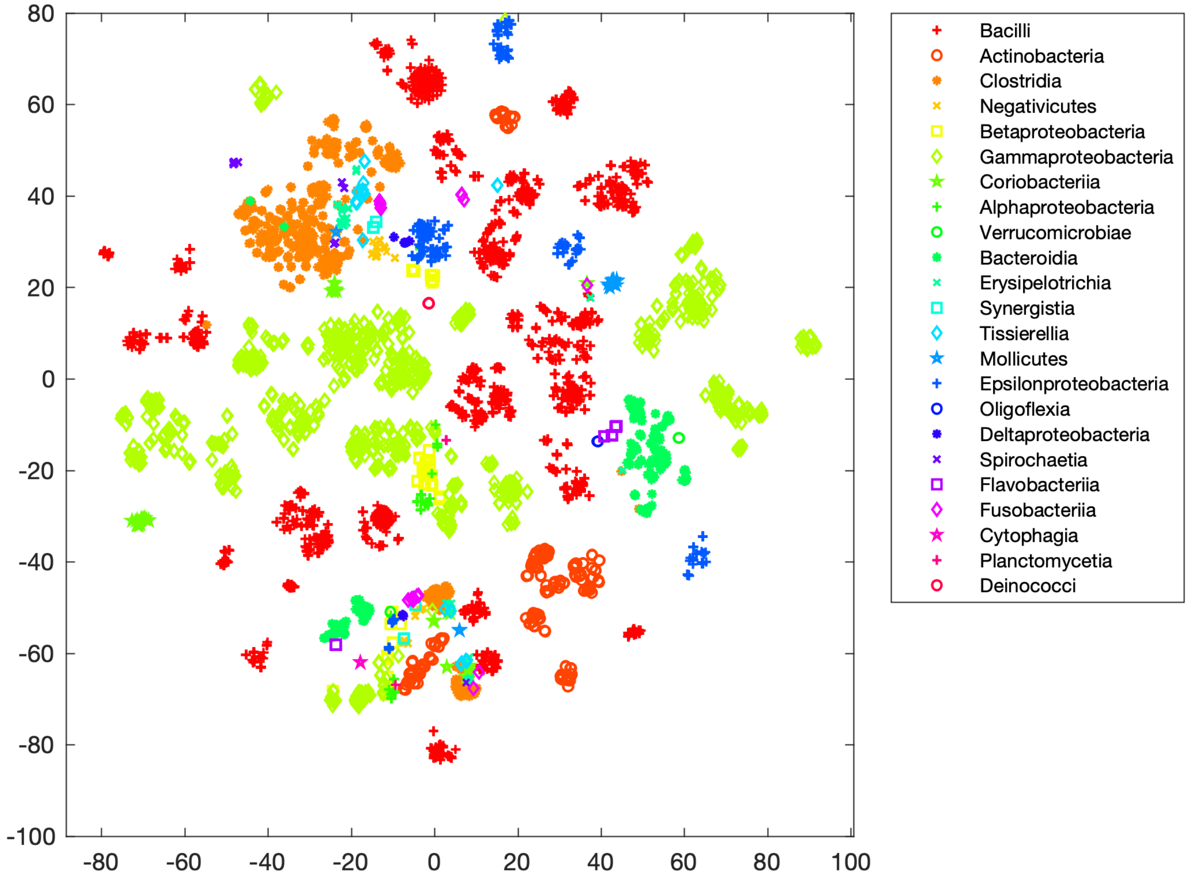


**Figure S3a)**: Clustering through t-distributed stochastic neighbour embedding (t-SNE) of reaction presence across all pathways per reconstruction for the 7,206 AGORA2 draft reconstructions retrieved from KBase. Shown are the members of the 23 largest classes by class.

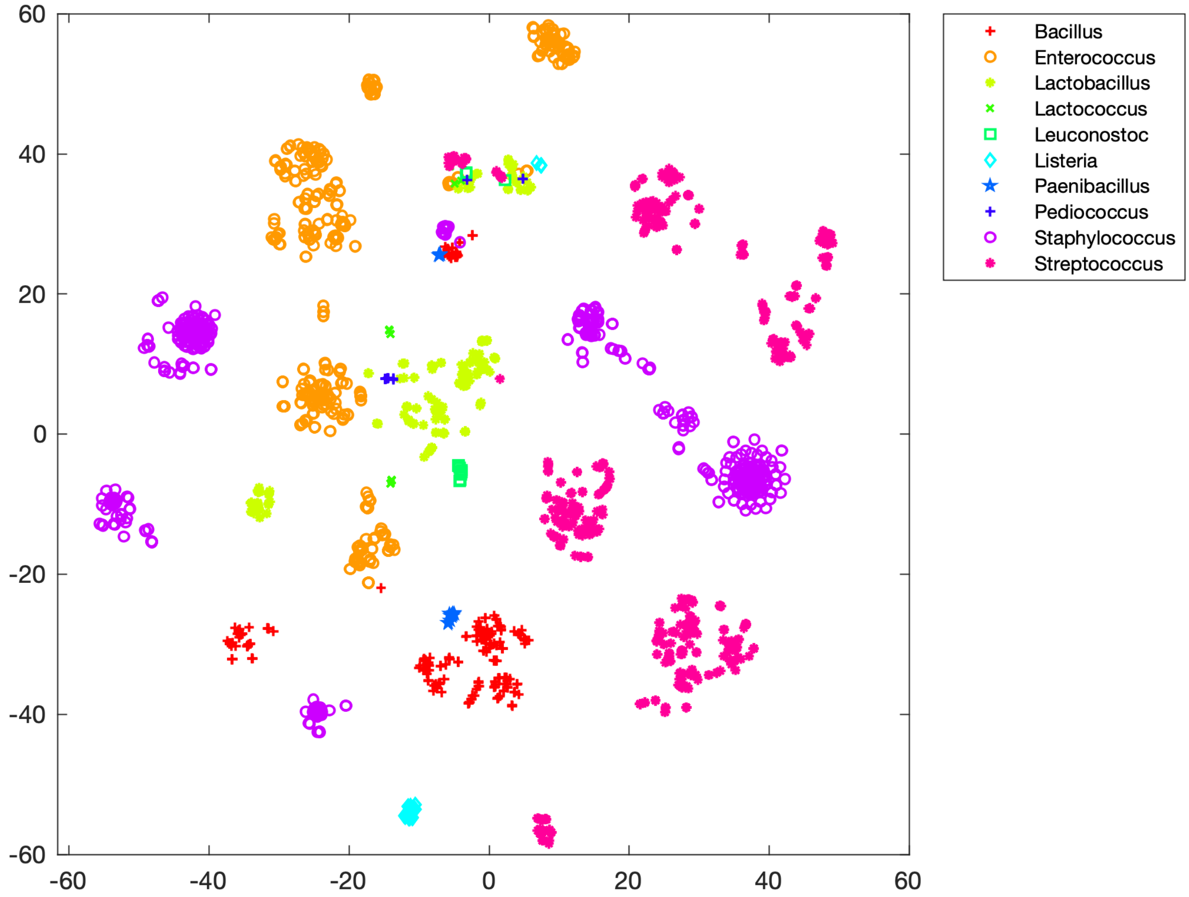


**Figure S3b)**: Clustering through t-distributed stochastic neighbour embedding (t-SNE) of reaction presence across all pathways per reconstruction for the 7,206 AGORA2 draft reconstructions retrieved from KBase. Shown are the members of the Bacilli class by genus.

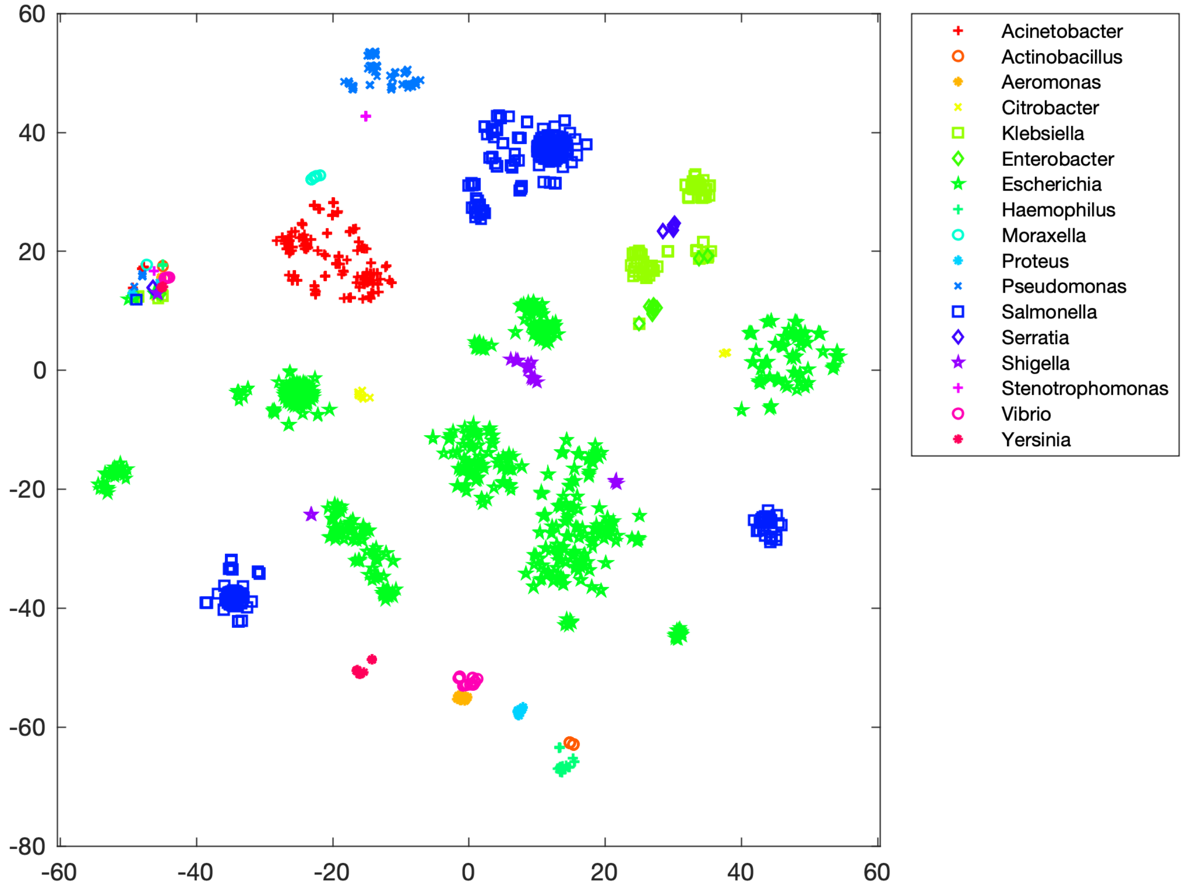


**Figure S3c)**: Clustering through t-distributed stochastic neighbour embedding (t-SNE) of reaction presence across all pathways per reconstruction for the 7,206 AGORA2 draft reconstructions retrieved from KBase. Shown are the members of the Gammaproteobacteria class by genus.

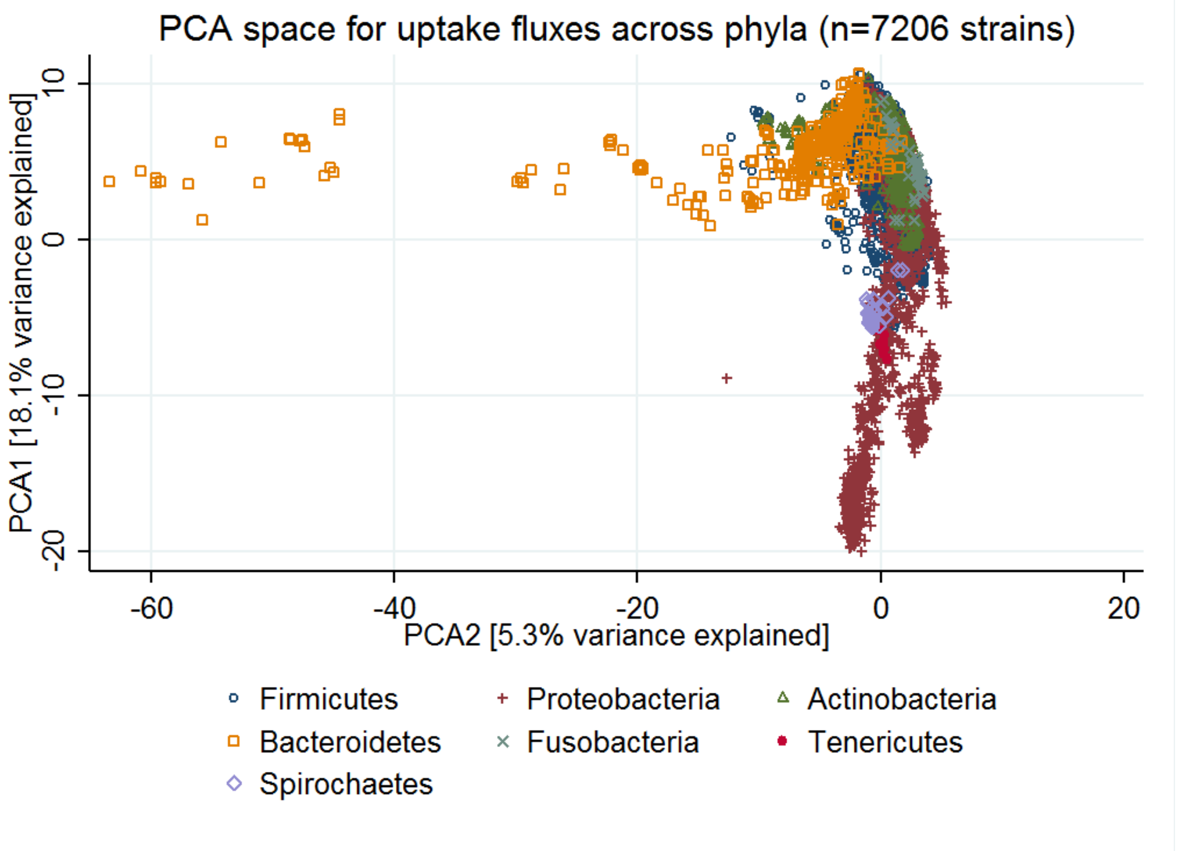


**Figure S4a**: Principles Component Analysis (PCA) space (first two dimensions) of the uptake fluxes under a European diet for 7206 strains. Displayed are the strains belonging to the seven biggest phyla in AGORA2 (Firmicutes, Proteobacteria, Actinobacteria, Bacteroidetes, Fusobacteria, Tenericutes, Spirochaetes).

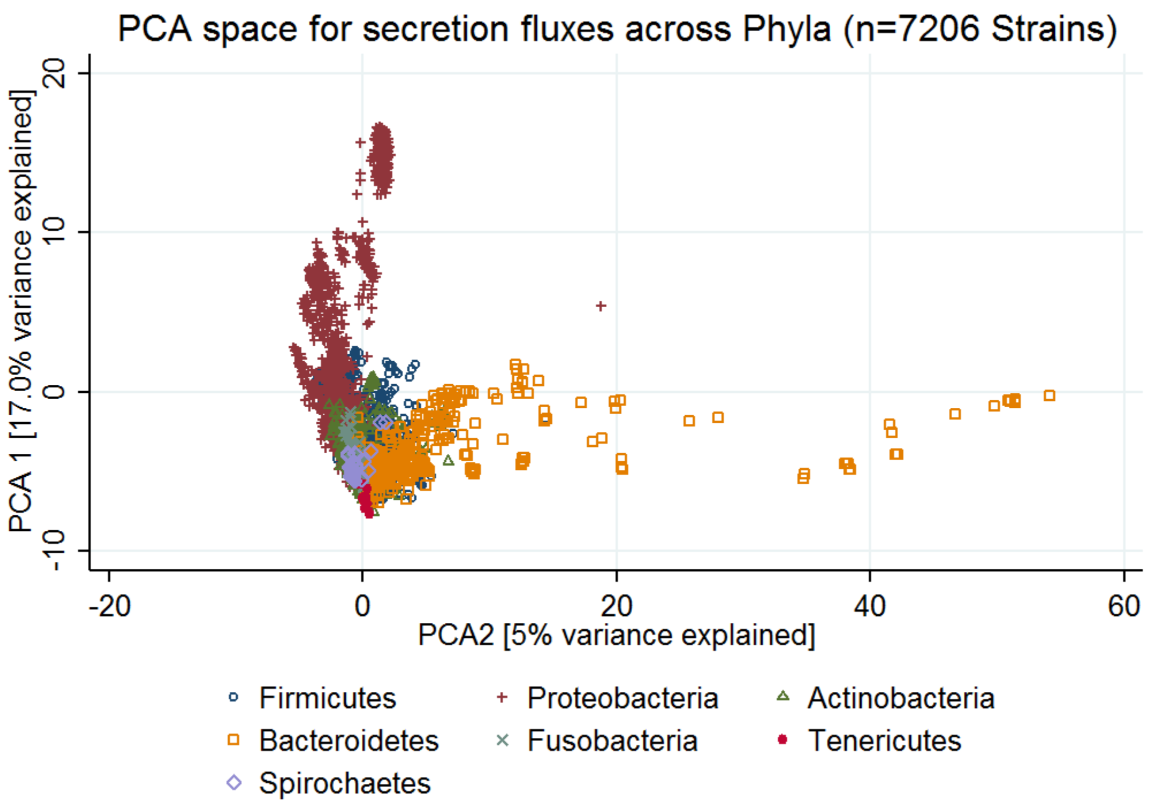


**Figure S4b**: Principles Component Analysis (PCA) space (first two dimensions) of the secretion fluxes under a European diet for 7206 strains. Displayed are the strains belonging to the seven largest phyla in AGORA2 (Firmicutes, Proteobacteria, Actinobacteria, Bacteroidetes, Fusobacteria, Tenericutes, Spirochaetes).

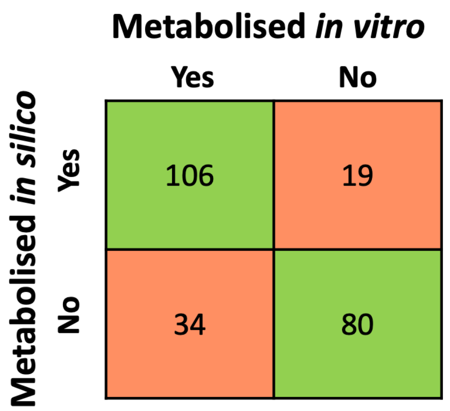


**Figure S5**: Overlap between independent, experimentally demonstrated activity of drug-metabolising enzymes and predictions by models derived from the AGORA2 reconstructions. 238 drug-microbe pairs Accuracy=78.2% (186 out of 238) with a sensitivity of 75.7% and a specificity of 80.8%. The models were grown on unlimited medium and flux through the appropriate reaction(s) for each drug conversion (Table S7) was tested. Accuracy=0.78, sensitivity= 0.76, and specificity= 0.81. The models were grown on unlimited medium and flux through the appropriate reaction(s) for each drug conversion (Table S7) was tested. Accuracy=0.78, sensitivity= 0.76, and specificity= 0.81.

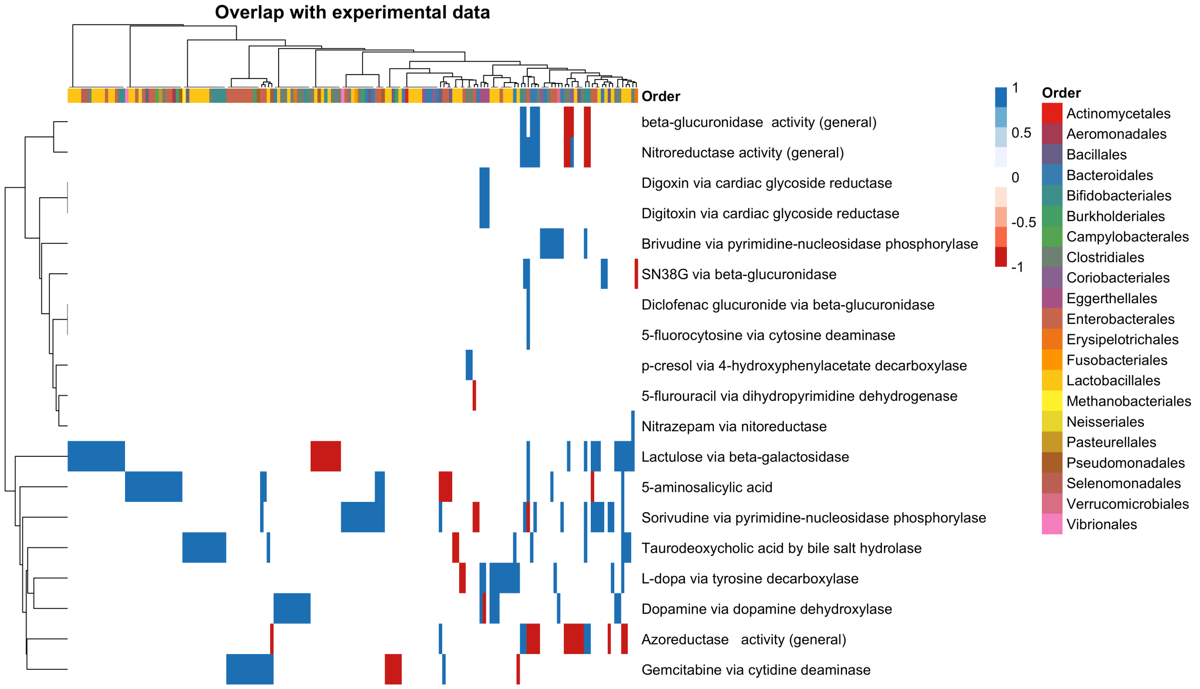


**Figure S6**: Overlap of AGORA2 model predictions for drug enzyme activity with published *in vitro* data. The experimental findings that the models were tested against are shown in Table S7. Columns represent strains/species (annotated with the respective order classifications) and rows represent drug-metabolising enzymes that were tested. Blue=model agrees with *in vitro* data, red=model disagrees with i*n vitro* data, white=no data available for this strain/species and enzyme.

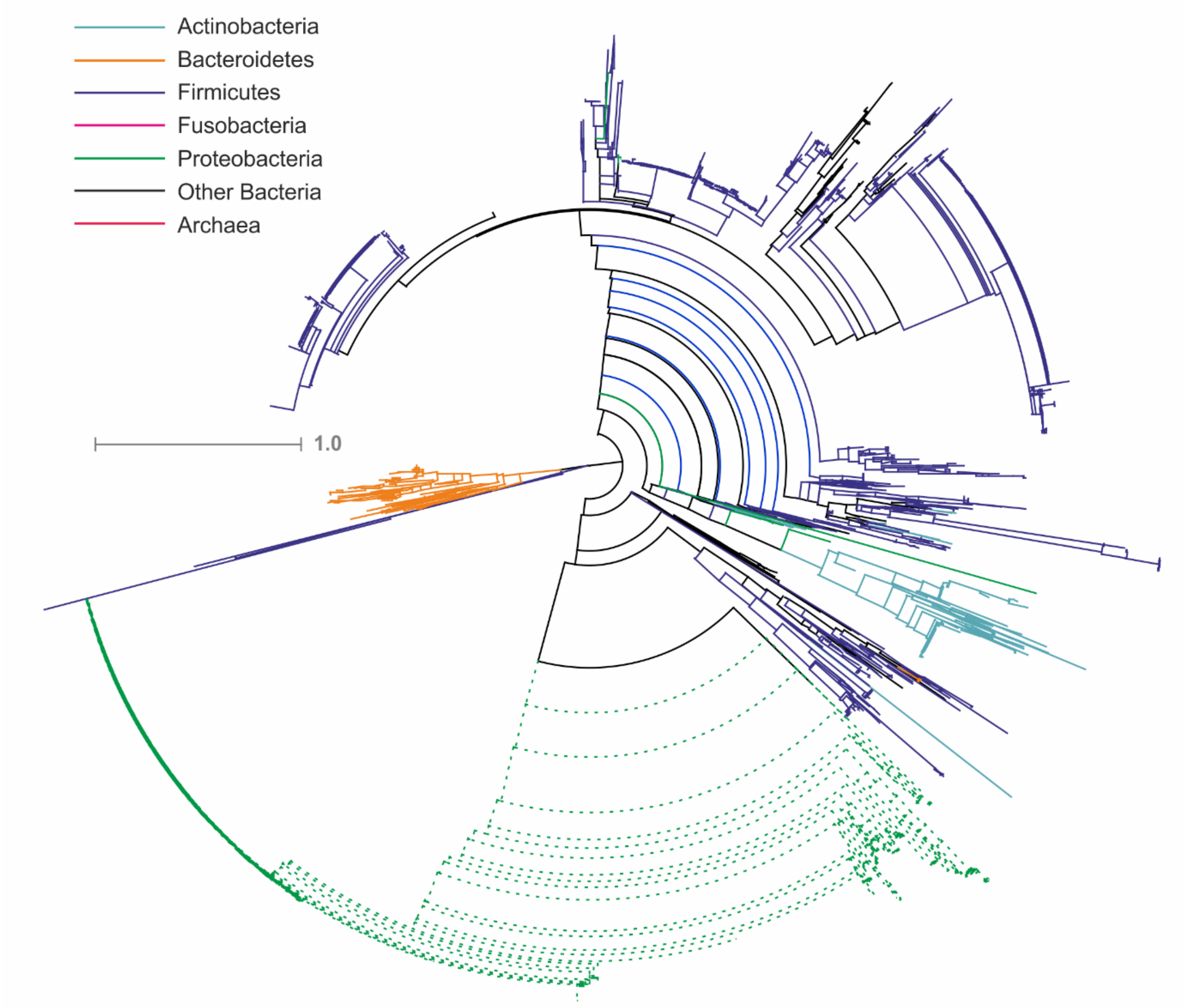


**Figure S7a:** Maximal-likelihood phylogenetic tree for cytidine deaminase (cCda, eCda, EC: 3.5.4.5) proteins in the analysed genomes. Taxonomy is shown by branch colour; solid lines, cytoplasmic proteins; dotted lines, extracellular / periplasmic proteins.

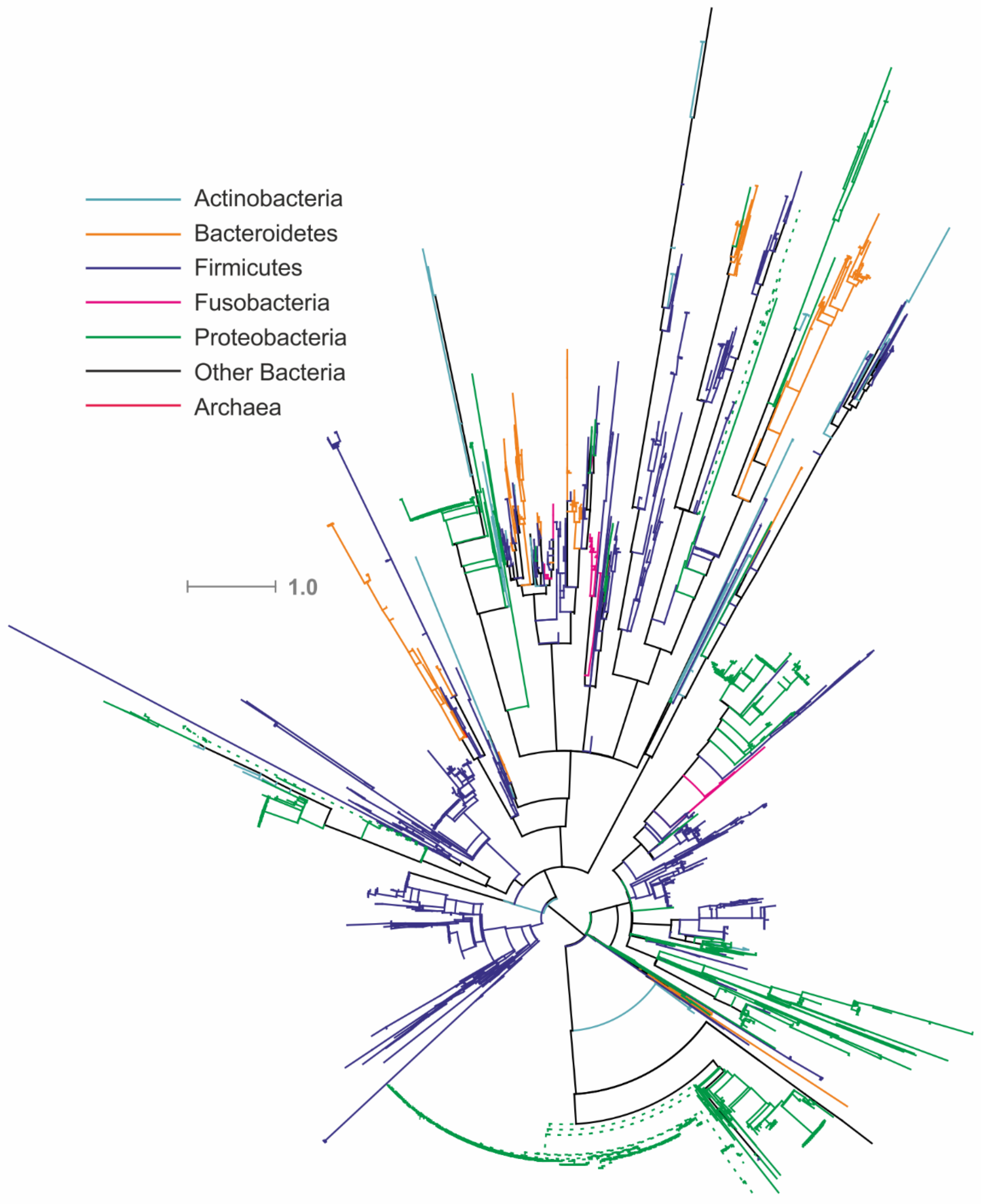


**Figure S7b:** Maximal-likelihood phylogenetic tree for nitroreductase (cNit, eNit, EC: 1.-.-.-) proteins in the analysed genomes. Taxonomy is shown by branch colour; solid lines, cytoplasmic proteins; dotted lines, extracellular / periplasmic proteins.

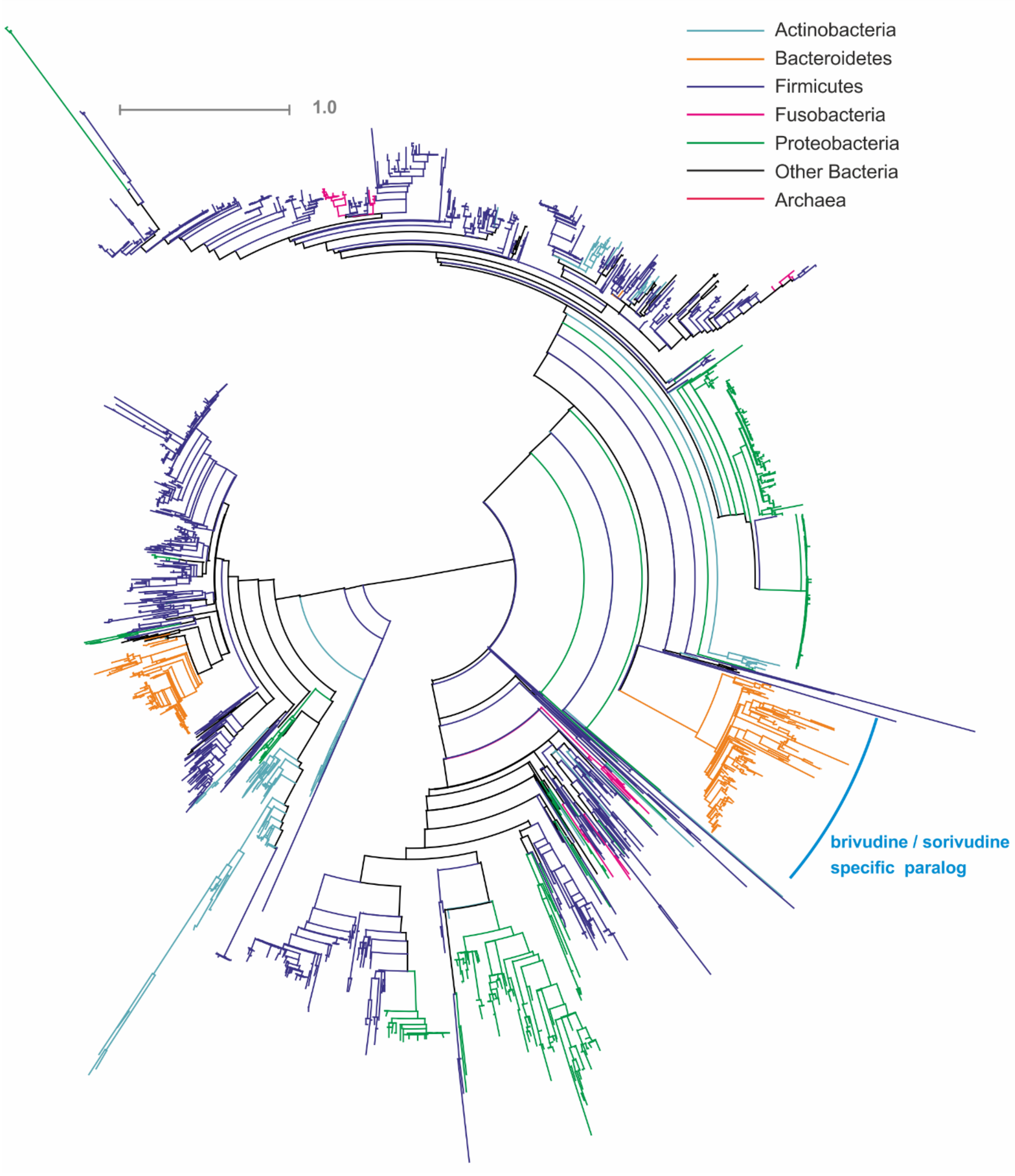


**Figure S7c:** Maximal-likelihood phylogenetic tree for pyrimidine-nucleoside phosphorylase (cBRV, EC: 2.4.2.2) proteins in the analysed genomes. Taxonomy is shown by branch colour.

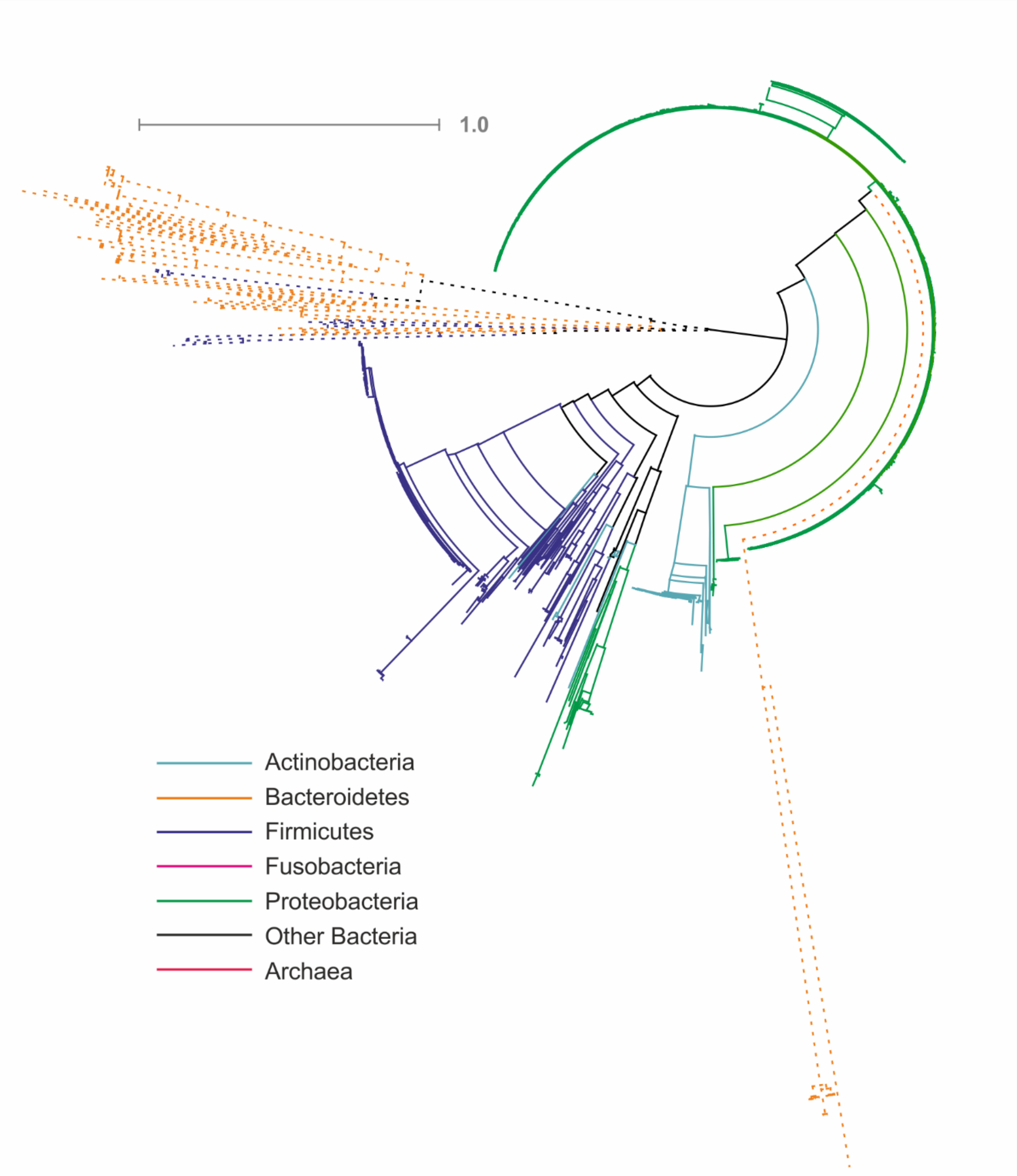


**Figure S7d:** Maximal-likelihood phylogenetic tree for β-glucuronidase (cUidA, eUidA, EC: 3.2.1.31) proteins in the analysed genomes. Taxonomy is shown by branch colour; solid lines, cytoplasmic proteins; dotted lines, extracellular / periplasmic proteins.

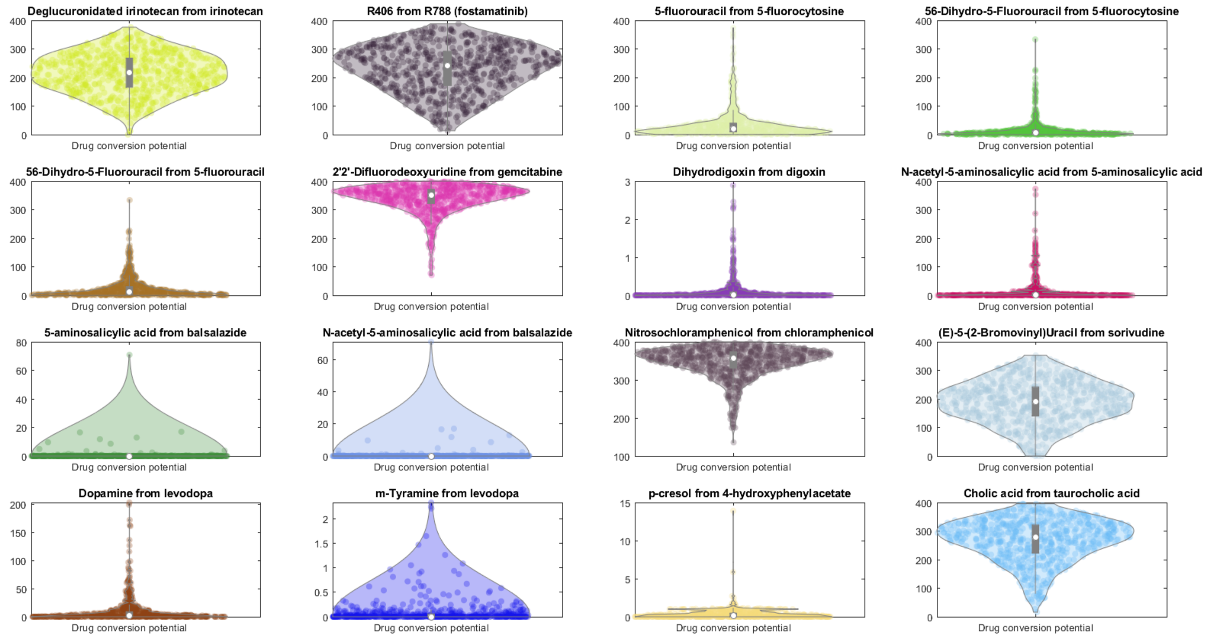


**Figure S8**: Drug conversion potential in the microbiomes of 616 Japanese colorectal cancer patients and controls on the Average Japanese diet plotted against the total abundance of the respective drug-metabolising enzymes. The violin plots show the distribution of drug metabolite flux in mmol/person/day. See Figure 3a for a description of each drug-metabolising enzyme.

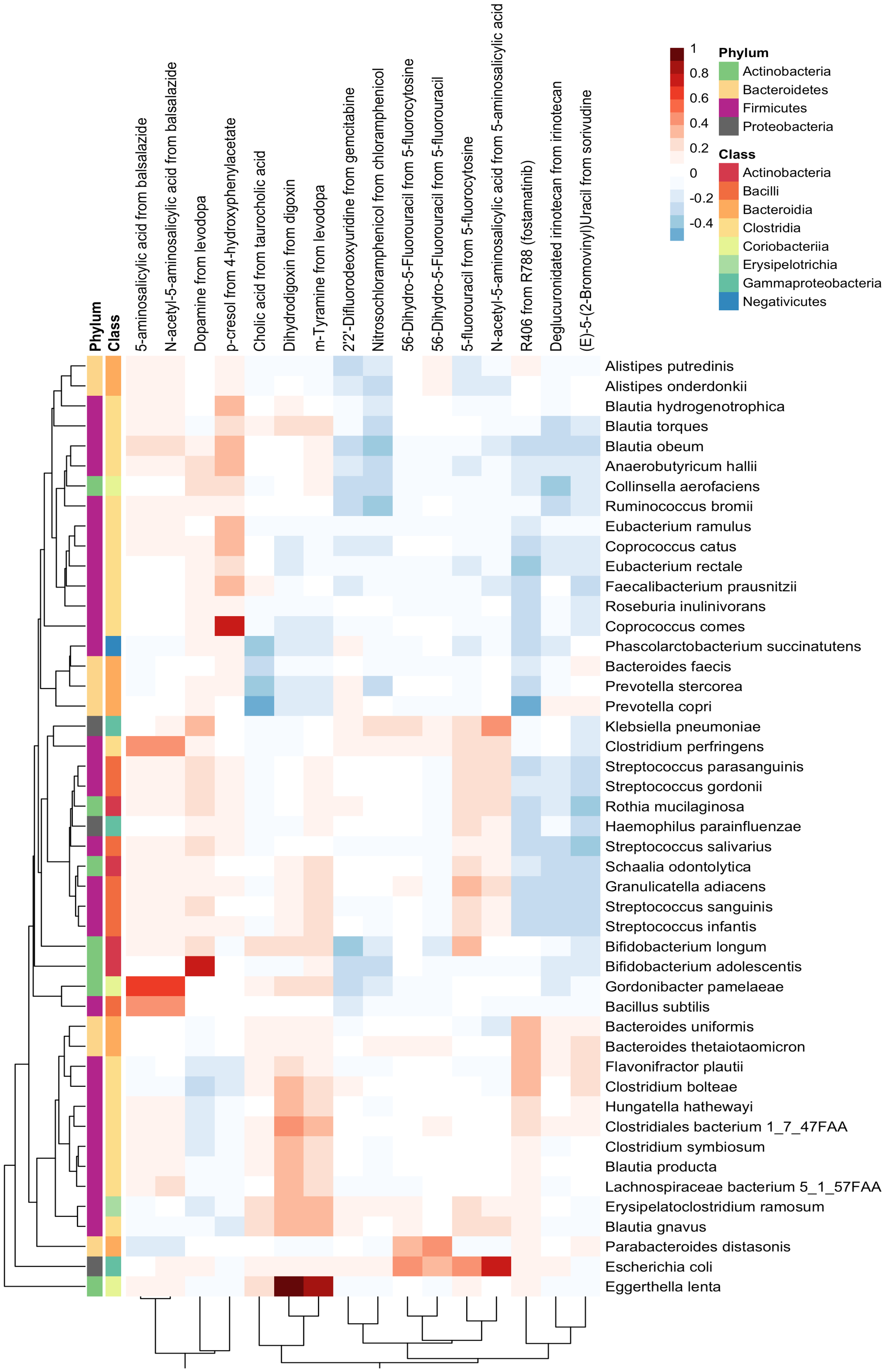


**Figure S9**: Spearman correlations between species abundances and drug conversion potential (mmol/person/day) in 616 microbiomes of Japanese colorectal cancer patients and controls.

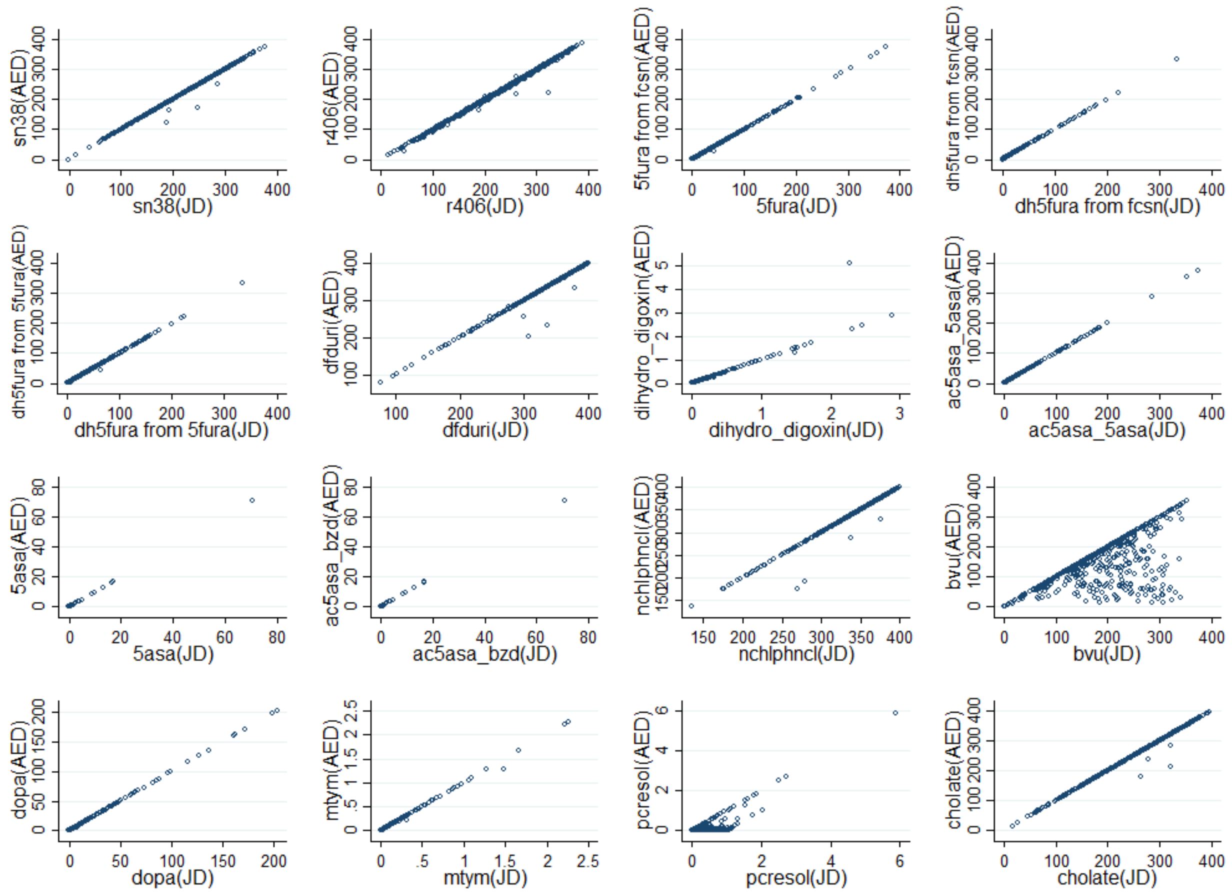


**Figure S10:** Scatter plots of drug-metabolising potentials under a European average diet (AED) against the metabolising potential under a Japanese diet (JP). Sn38=deglucuronated irinotecan, 5fura=5-fluorouracil, fcsn=5-fluorocytosine, dh5fura=5,6.dihydro-5-fluorouracil, dihydro_digoxin=Dihydrodigoxin, ac5asa=N-acetyl-5-aminosalicylic acid, 5asa=5-aminosalicylic acid, dfduri=2',2'-Difuorodeoyuridine, ac5asa_bzd=N-acetyl-5-aminosalicylic acid from balsalizide, nchlphncl=Nitrosochoramphenicol, bvu=(E)-5-(2-Bromovinyl)Uracil, dopa=Dopamine, mtym=m-tyramine, cholate=cholic acid.

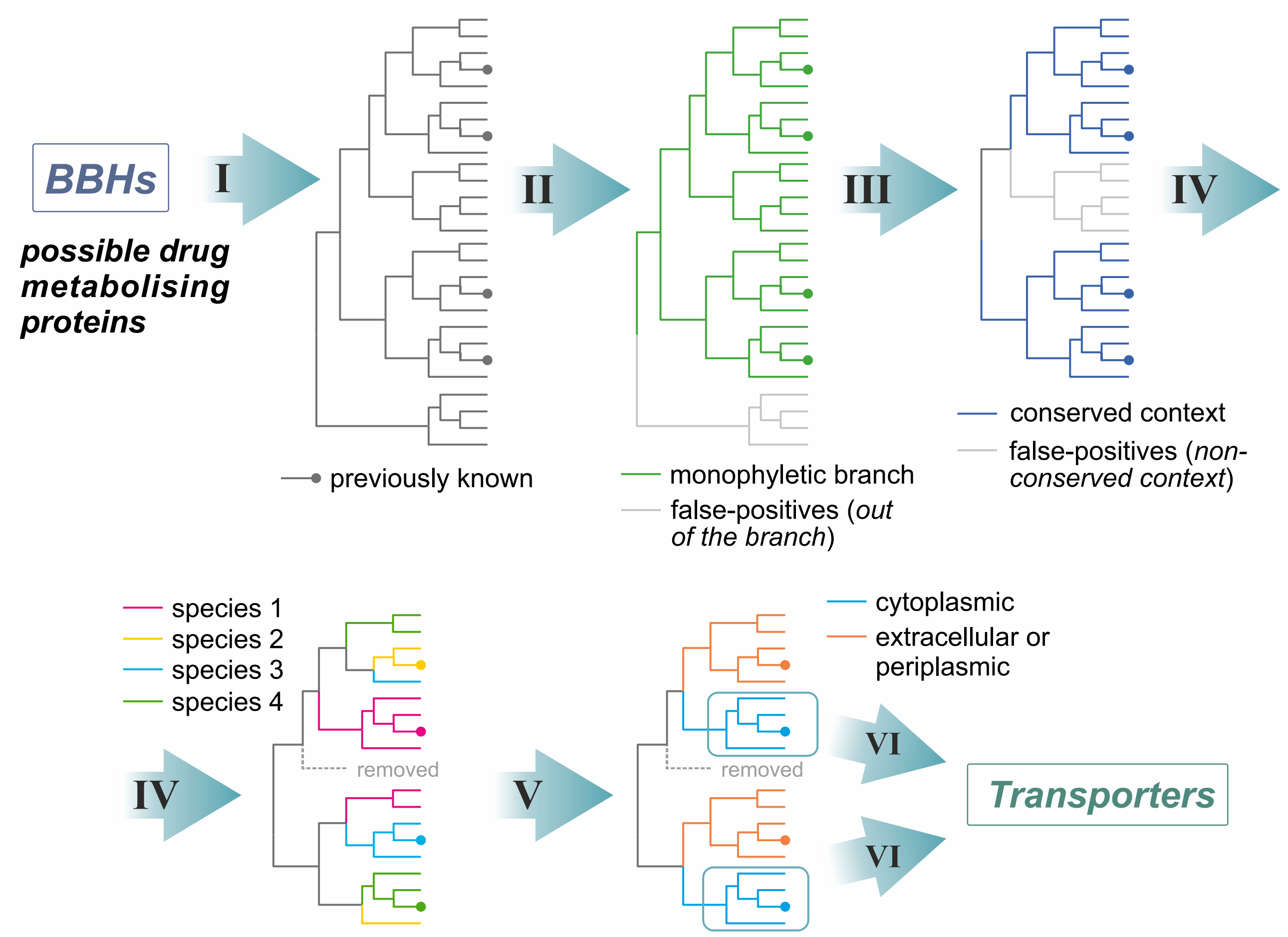


**Figure S11:** Procedures for a manual refinement of the drug-metabolising genes. The following steps are shown: (I) construction of the maximal-likelihood phylogenetic tree, rooted at a mid-point, and mapping of the previously known proteins; (II) defining monophyletic branches including all the previously known proteins; (III) removal of the false-positive predictions and analysis of the genomic context; (IV) removal of the false-positive predictions and defining of the species-specific protein clusters; (V) prediction of the subcellular localisation; (VI) analysis of the genomic context to predict transporters, only for cytoplasmic enzymes.
